# Phenotype Driven Data Augmentation Methods for Transcriptomic Data

**DOI:** 10.1101/2023.10.26.564263

**Authors:** Nikita Janakarajan, Mara Graziani, María Rodríguez Martínez

## Abstract

With machine learning taking over biomedical applications, working with transcriptomic data on supervised learning tasks is challenging due to high dimensionality, low patient numbers and class imbalances. Machine learning models tend to overfit these data and do not generalise well on out-of-distribution samples. Data augmentation strategies help alleviate this by introducing synthetic data points and acting as regularisers. However, existing approaches are either computationally intensive, require population parametric estimates or generate insufficiently diverse samples. To address these challenges, we introduce two classes of phenotype driven data augmentation approaches – signature-dependent and signature-independent. The signature-dependent methods assume the existence of distinct gene signatures describing some phenotype and are simple, non-parametric, and novel data augmentation methods. The signature-independent methods are a modification of the established Gamma-Poisson and Poisson sampling methods for gene expression data. As case studies, we apply our augmentation methods to transcriptomic data of colorectal and breast cancer. Through discriminative and generative experiments with external validation, we show that our methods improve patient stratification by 5 − 15% over other augmentation methods in different cases. The study additionally provides insights into the limited benefits of over-augmenting data. The code is hosted on <u>GitHub</u>, and includes a link to the augmented datasets.

## 1 Introduction

The application of machine learning in biomedicine, particularly for supervised learning tasks pertaining to cancer, is becoming increasingly popular. Technological advances has provided increasing access to transcriptomics data, catalysing an increase in research aimed at pattern discovery, phenotype classification, disease subtyping, survival analysis, and more [1]. However, inherent challenges in this domain persist, notably the high dimensionality of the data and the limited number of samples [2]. This is particularly problematic for supervised learning tasks because it is often the case that the class labels are imbalanced in such datasets [3]. All of these challenges combined make machine learning models prone to over-fitting and reduces their ability to generalize to new datasets [4], even for the same samples measured at different locations [5].

Data augmentation methods are typically used to alleviate these issues with limited sample numbers. These methods generate new data points from existing ones, addressing any class imbalances and under-representations. The most commonly used method is random oversampling or sampling with replacement [6]. While effective for constructing accurate, less biased classifiers [7], this method, however, does not generate new samples and so the model gains no new information about the underlying distribution. This affects model generalisation and robustness to unseen datasets. Although new sample generation can be achieved by a weighted mixing of observations [8], assigning unique class labels to such samples can be challenging, making it less suitable for supervised learning. Oversampling by interpolation is another popular method of augmenting biological data. Synthetic Minority Oversampling Technique (SMOTE) [9] is a prime example of this sampling strategy, achieving state of the art performances on tasks affected by class imbalances. However, the method comes with its challenges: (i) for high-dimensional datasets, SMOTE does not attenuate the bias towards the majority class [10], (ii) when applied to high-dimensional data, SMOTE preserves the expected value of the minority class while decreasing its biological variability, which can be problematic for classifiers that rely on class-specific variances [10], (iii) SMOTE can be computationally demanding due to extensive pairwise distance calculations.

Data can also be augmented using methods that rely on population statistics and data models. In the case of transcriptomics data, this translates into using parametric methods such as the Poisson distribution [11–13] or the negative binomial distribution, also known as the Gamma-Poisson distribution [14, 15] to sample new data. To solve class imbalance, these methods require class-specific parameter estimations. Challenges arise when the data distribution support is too large, leading to low data density in some regions and potentially generating uninformative data distributions. Conversely, limited support can result in insufficient sample diversity. Furthermore, out-of-distribution data points can also impact parameter inference, making these methods sensitive to outliers and measurement errors [16]. Deep learning based methods, such as Generative adversarial network (GAN) [17] and its variants, instead learn the distribution parameters to create synthetic gene expression samples [8, 18, 19]. However, these methods are computationally intensive, difficult to train, require detailed knowledge about gene regulatory networks and a large amount of data, which is difficult to obtain, particularly in cancer studies.

Given the challenges faced by current approaches in generating new samples for supervised learning, we propose two types of phenotype or target driven approaches for data augmentation. The first type leverages phenotypic gene signatures – sets of genes with distinct expression patterns that hold prognostic, diagnostic or predictive value – to drive the augmentation. Gene signatures are invaluable for phenotypic predictions, and are especially useful in cancer subtyping [20–22]. In a nutshell, patients’ gene signatures within and between phenotype variants are mixed to generate new observations. This set of methods is computationally efficient and does not require estimating distribution parameters. The second type is a modification of the parametric methods - Gamma-Poisson and Poisson sampling - for cases where signatures are not known or entirely distinct for a phenotype. To address challenges with class-imbalance, we adapt these methods to initialise a new distribution based on a subset of the data to generate a new sample.

We demonstrate the effectiveness of our sampling methods on cancer-related tasks. We believe limitations in data availability is a contributor to its status as one of the least understood diseases. Moreover, as significant work has been done in the field to identify associated signatures, it presents us with a playground for testing signature-dependent and -independent methods particularly in sub-type identification. We make use of colorectal (TCGA COADREAD (Colon Adenocarcinoma and Rectal Adenocarcinoma)) and breast cancer (GSE20713, TCGA BRCA (Breast Cancer)) gene expression datasets for training and evaluate the generalisation performance on the out-of-domain CPTAC COAD (Colon Adenocarcinoma) and METABRIC datasets, respectively. Our methods successfully address class imbalances, improve model performance, and offer a promising solution for data augmentation in the biomedical domain. We envision our methods to be applicable in modeling mixed phenotype scenarios, particularly useful for cancer heterogeneity studies, and in developing models that are robust against class imbalance, a problem that plagues the biology domain.

Our contributions are as follows:

1. We introduce a novel, non-parametric signatue-dependent data augmentation method and benchmark it against conventional techniques;
2. We demonstrate the utility of the Gamma-Poisson and Poisson distributions as class balancing signature-independent data augmentation strategies by proposing a modification in the sampling strategy;
3. We highlight the influence of augmentation size on model performance through a series of experiments in both discriminative and generative settings.

## 2 Methods

### 2.1 Signature-dependent sampling

Signature-driven sampling aims to create new data points from existing data points while increasing diversity in the generated samples. Simply put, to generate new samples for a given phenotype, we leverage the high informativeness of the gene signatures associated to that particular phenotype, while assuming that the signatures of the other phenotypes in that sample are less informative for the given phenotype. We investigate two ways of augmenting the data samples:

1. *Intra-class crossover:* Crossing over between samples having the same phenotypic variant.
2. *Inter-class crossover:* Crossing over between samples across all phenotypic variants.

In the following sections, we describe the method through a practical example on the gene expression of patients with colorectal cancer, using the Consensus Molecular Subtype (CMS) as phenotype. This phenotype has four variants, namely, CMS1, CMS2, CMS3, and CMS4 [23], and each CMS class has a set of 10 genes that are highly predictive of it [24]. This greatly reduces the dimensionality of our gene expression dataset and gives us a panel of 40 genes that can be used for classifying patients into the 4 colorectal cancer subtypes. We describe the two methods of crossing over using this dataset as our premise. Formal definitions of the methods can be found in Appendix A and B, respectively.

#### 2.1.1. Intra-class crossover sampling

To limit the likelihood of mixing phenotypes, and hence introducing samples of dubious labels, we first perform crossing over between samples belonging to the same phenotype. Thus, in this setting, new samples are generated only from samples having the same phenotype and so naturally, the synthetic sample inherits that phenotype. The crossing-over between samples is done at the gene signature level.

Figure 1C illustrates the process of intra-class crossover sampling. Consider an ordered gene matrix where the patients and the genes are grouped according to their CMS class. To generate a new CMS1 sample, we randomly sample each of the four gene signature blocks from the subset of all CMS1 patients only.

**Figure 1:**
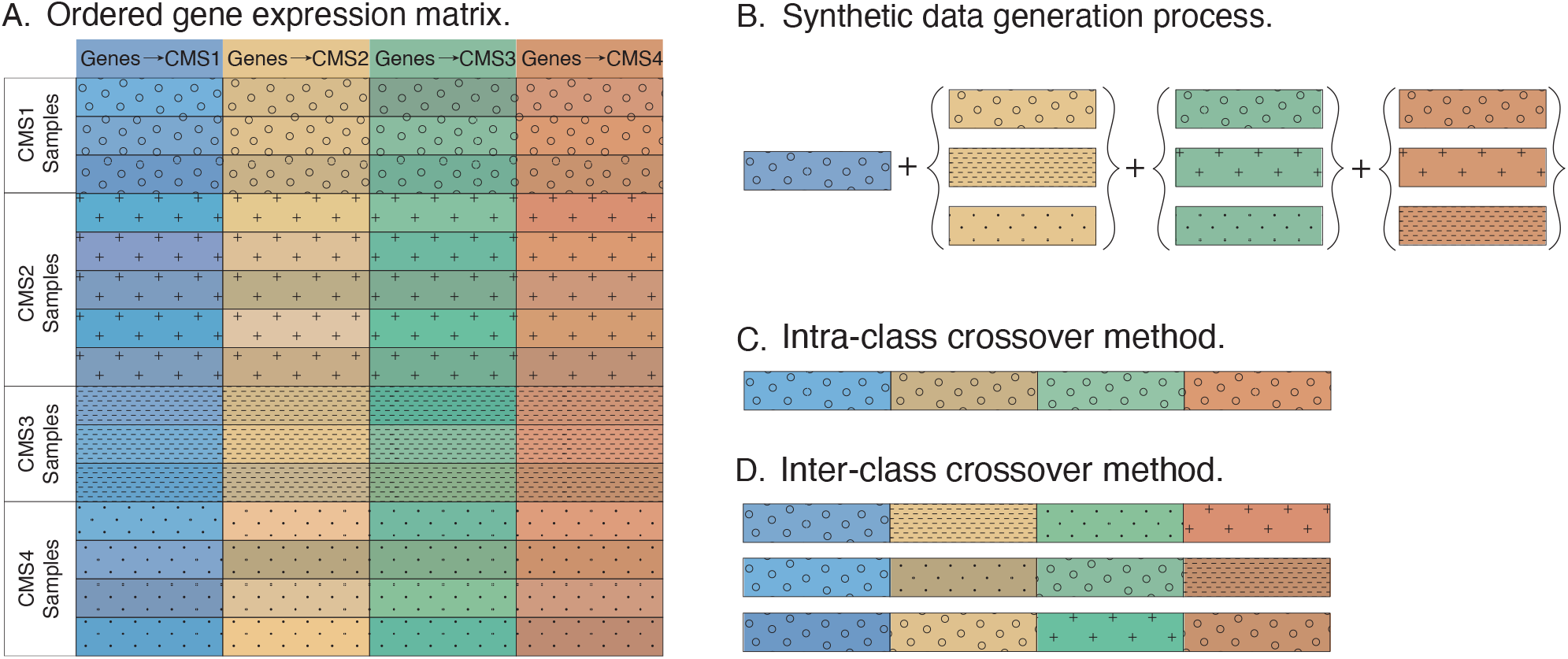
Crossover sampling strategy. A. Consider a gene expression matrix where each gene block, identified by colour, is the predictive signature of a phenotype (CMS (colorectal cancer subtype)), identified by pattern. The rows represent patients with a given phenotype, and the shade of the gene block indicates a patient-dependent expression pattern. B. To generate a new sample, we mix and match gene signature blocks under certain phenotypic constraints. C. Intra-class crossover sampling. A sample of class CMS1 is generated by sampling gene signature blocks from the subset of CMS1 patients only. D. Inter-class crossover sampling. A sample of class CMS1 is generated by sampling the CMS1 gene signature block from the subset of CMS1 patients only and each of the remaining gene blocks are sampled from patients who do not belong to the CMS class that the gene block predicts.

Since we sample the blocks only from the subset of CMS1 patients, the expression values of genes associated with the other CMS types are typical of what is observed in a CMS1 patient. This process is repeated for the other CMS types, that is, a CMS2 patient is sampled only from the subset of patients belonging to CMS2, and so on. Restricting the sampling subset also allows this method to handle any correlations between genes in a secure way. A more formal definition of the method is described in Appendix A.

#### 2.1.2. Inter-class crossover sampling

In Inter-class crossover sampling, signature blocks are sampled from all samples. To avoid ambiguous labels, we add a constraint such that the defining signature block can only come from a sample that belongs to the phenotype being sampled. This method uses the entire dataset to create new samples, in contrast to intra-class crossover sampling, which samples from a phenotype-specific subset of the data. A detailed description of the method can be found in Appendix B.

Inter-class crossover sampling is illustrated in Figure 1D in the context of CMS classes. When creating a CMS1 sample, we first sample the defining signature block from the set of all CMS1 patients. This ensures that the predictive signature block has typical values of a CMS1 patient. The remaining gene blocks are sampled from the subset of all patients, excluding those patients that belong to the class associated with the gene block being sampled. This prevents any ambiguity in class label assignment. For example, the CMS2 block can be sampled from all patients except those belonging to CMS2 since the expression values of the CMS2 block in the other patients are not typically predictive of CMS2.

### 2.2 Signature-independent sampling

Parametric methods such as the Gamma-Poisson and Poisson distributions are popular augmentation methods for RNA-Seq data. In our experiments, we modify these two methods to improve the class-specific diversity of the generated samples over the original versions. Briefly, the distributional parameters are estimated from randomly sampled sub-populations for every new sample that is

#### 2.2.1 Modified Gamma-Poisson sampling

The negative-binomial (NB) distribution is a commonly used parametric method for augmenting RNA-Seq data. The negative-binomial (NB) distribution is typically expressed as a Gamma-Poisson mixture [25, 26] to make it more tractable. In this formulation, the rate parameter *λ* of the Poisson distribution is a random variable sampled from a gamma distribution parameterised by the shape (*α*) and rate (*β*) parameters, estimated from the observed population. Our proposed modification to the sampling strategy is to create a mixture of Gamma-Poisson distributions. To generate every new observation, a subset *S* of samples is randomly chosen. The mean *μ* and variance *σ*^2^ of this subset *S* is used to estimate the *α* and *β* parameters, which initialise the gamma distribution as shown in Equation 1. Since the augmentation is aimed at solving class imbalance, the subset is created from samples belonging to the same class. Thus, the newly generated observation will have the same label as the subset samples. The size of the subset |*S*| is a hyperparameter defined by the user. A random variable is then sampled from this gamma distribution and is used to initialise the Poisson distribution, from which a new observation is sampled. This process is repeated *n* times to generate *n* new observations. A more detailed explanation is described in Appendix C.

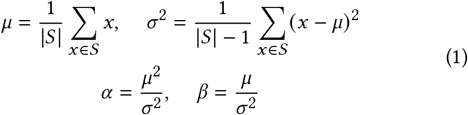

#### 2.2.2 Modified Poisson sampling

Similarly to the Gamma-Poisson strategy, a subset is first sampled from the class to be augmented, after which the mean is computed and defined as the rate parameter *λ* for the Poisson distribution as shown in Equation 2. A new observation is sampled from this distribution and this process is repeated *n* times to generate *n* new observations. A more detailed explanation is described in Appendix D.

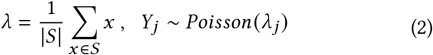

## 3 Experimental Setting

### 3.1 Datasets

We use TCGA COADREAD [27] and CPTAC [28] RNASeq datasets for the first part of our study demonstrating the signature dependent methods. TCGA COADREAD is designated for training and in-domain testing, while the CPTAC COAD gene expression dataset is used exclusively for out-of-domain validation. As an initial preprocessing step, we filter out replicates and only consider patients with primary tumours and available phenotype labels. The genes are reduced to the signature genes associated with these phenotypes. For the second part of our study addressing edge cases such as unavailable, overlapping or correlated signatures, we utilise the GSE20713 microarray breast cancer data [29] for training and in-domain testing, and METABRIC [30, 31] dataset for out-of-domain validation. Summary of class counts of all datasets are shown in Appendix Table 2 and Table 5, respectively.

### 3.2 Data augmentation process

Data augmentation is applied only to training data. The external datasets CPTAC and METABRIC used for validation are not augmented. The following augmentation methods are benchmarked - (i) Inter-class crossover sampling, (ii) Intra-class crossover sampling, (iii) Modified Gamma-Poisson sampling (Mod. GP), (iv) Modified Poisson sampling (Mod. Poisson), (v) Synthetic Minority Oversampling Technique (SMOTE) [32], (vi) Random oversampling (Replacement), and (vii) No augmentation (Unaugmented). The train and in-domain test splits are created from the TCGA COADREAD count and GSE20713 RMA-normalised microarray training datasets.

In the case of TCGA COADREAD, we consider 3 class sizes for augmentation - ‘Max’, 500 and 5000. In the ‘Max’ setting, only the minority classes are oversampled to match the size of the majority class. For class sizes 500 and 5000, all classes are augmented until the number of samples in each class reaches 500 and 5000, respectively. In addition to testing the generalisability of the augmentation methods, the different sizes also provide insights into the effect of size on the quality of the augmented data. For the unaugmented case, we simply take the log-FPKM normalised expression data of TCGA COADREAD. A visualisation of the augmented TCGA COADREAD data is provided in Appendix E. Details on hyperparameter selection for Mod. Gamma-Poisson and Mod. Poisson are described in Appendix F. After augmenting TCGA COADREAD count data, we perform FPKM normalisation (Fragment Per Kilobase per Million mapped reads) and log transformation as per [33], based on the training split.

Since GSE20713 microarray data are one order of magnitude smaller than TCGA COADREAD, augmentations are also one order of magnitude smaller, that is, we augment the data to class sizes ‘Max’, 50 and 500. For the unaugmented case, we simply take the RMA-normalised expression data as provided. Since microarray data from GSE20713 is already RMA-normalised, we do not do any further processing.

### 3.3 Models

For all discriminative tasks, we consider 5 commonly used classifiers to get an overall performance estimate, namely, Logistic Regression (LR), K-Nearest Neighbours (KNN), Support Vector Machines with RBF kernel (SVM-RBF), Explainable Boosting Machines (EBM), and Random Forests (RF) as implemented by sklearn. For the generative tasks, we use a variational autoencoder [34], implemented using PyTorch.

### 3.4 Evaluation criteria

We report the balanced accuracy (Appendix Equation 7) for all classification tasks on the in-domain and out-of-domain test sets. Balanced accuracy, as implemented by sklearn, is the average per-class recall (or sensitivity) scores. If the support samples for each class exhibit sufficient and meaningful diversity, the test classification performance on the augmented data is expected to outperform that of the unaugmented dataset. We perform a 5-fold stratified cross validation repeated 5 times, resulting in 25 unique training-test splits for each augmentation method and augmentation size. All significance tests are done with Wilcoxon signed-rank test adjusted to control the false discovery rate using the Benjamini-Hochberg method.

## 4 Results

In the first part of the experiments, we evaluate the generalisation performance of our proposed methods in two cases, (a) colorectal cancer subtype prediction (CMS) with non-overlapping signatures and (b) breast cancer PAM50 subtype prediction with overlapping signatures. In the second part of our experiments, we conduct an in-depth analysis of our methods using the colorectal cancer dataset. Implementation details of all experiments are described in Appendix G.

### 4.1 Generalisation performance

#### 4.1.1 Inter-class crossover sampling outperforms other methods when signatures do not overlap

We use colorectal cancer subtype prediction (CMS) to demonstrate the utility of our augmentation methods in the non-overlapping signature setting. We compare the generalisation performance in terms of balanced accuracy of the different augmentation methods in predicting the CMS class across the three class sizes - ‘Max’, 500 and 5000. The distribution of balanced accuracy on the unseen, out-of-domain CPTAC dataset across all classifiers and splits of data is illustrated in Figure 2. The inter-class crossover method generalises significantly better than the other augmentation methods. Particularly, inter-class crossover reports the highest overall performance for class sizes 500 (65.34% ± 0.032) and 5000 (66.45% ± 0.019), resulting in a 5% − 6% increased accuracy over unaugmented data (average balanced accuracy 59.9% ± 0.0297). The method also shows a 5% − 8% increased accuracy over data augmented by SMOTE (59.28% ± 0.0328 for class size 500 and 58.51% ± 0.0349 for class size 5000), and random oversampling (58.93% ± 0.0313 for class size 500 and 59.18% ± 0.0358 for class size 5000). We also observe that over-augmenting the data to class size 5000 brings little benefit in that only the variance across the splits is reduced. However, the difference in average performance is seemingly small compared to class size 500, suggesting that over-augmentation offers little to no benefits at the expense of increased computation.

**Figure 2:**
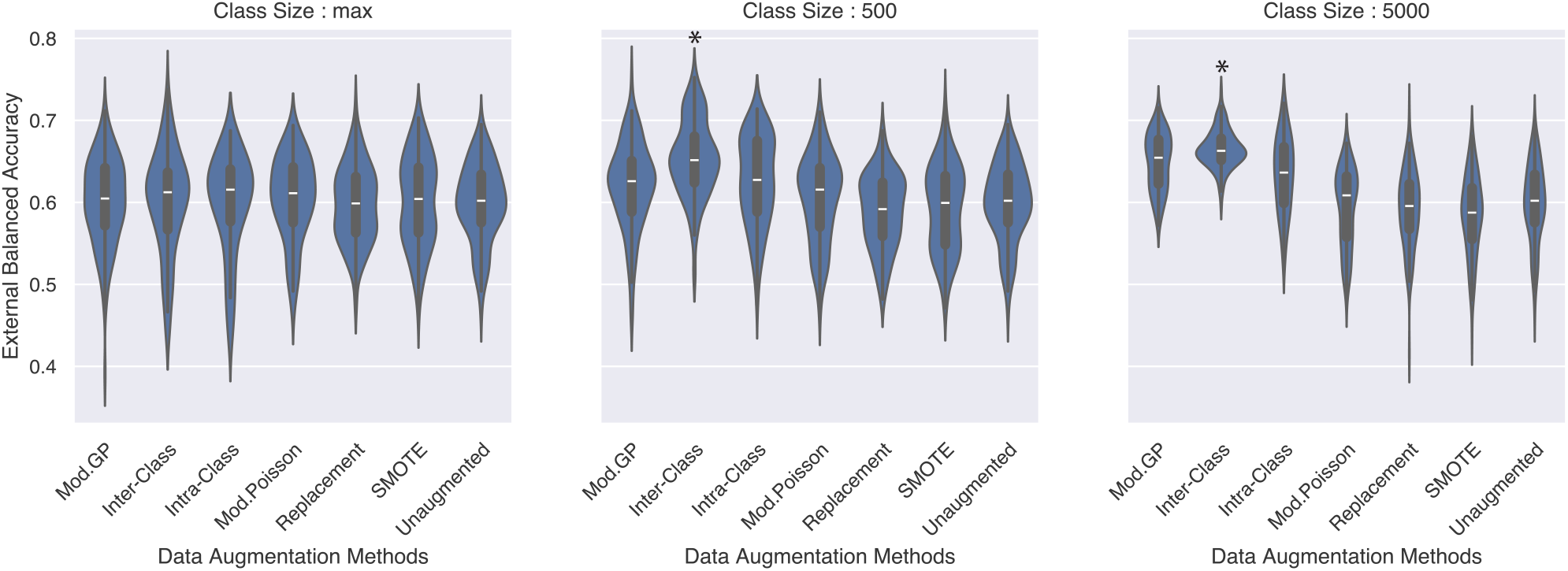
Distribution of balanced accuracy on out-of-domain CPTAC test set across all classification models (EBM, KNN, Logistic, RF, SVM-RBF) for each augmentation method, class size and CV split. The trend indicates that inter-class crossover method achieves the best generalisation performance and is a good candidate for tasks needing generalisation as it can achieve the best accuracy with fewer augmented samples compared to Mod. Gamma-Poisson. * indicates that the method is significantly different than the other methods.

Average balanced accuracy of each classification model, augmentation method, and class size is quantified in Appendix Table 10 for in-domain, and Appendix Table 11 for out-of-domain test sets. Additionally, the ROC-AUC scores are illustrated with a violin plot for both in-domain (Appendix Figure 12) and out-of-domain (Appendix Figure 13) test sets.

#### 4.1.2 Mod. Gamma-Poisson sampling outperforms other methods when signatures overlap

Although having signatures that are non-overlapping is ideal, it is often not the case. The signature-dependent methods cannot inherently handle overlapping genes as these methods rely on mixing blocks of genes. Overlapping genes complicate this procedure as the class being sampled and the relationships between these genes and classes need to be taken into account. The signature-independent methods, however, face no such challenge. Since sub-populations of the same class are sampled to initialise the sampling distributions, the inherent relationships between genes are maintained.

To demonstrate the limitations and strengths of these two approaches, we define a classification problem based on the PAM50 breast cancer subtypes - Luminal A, Luminal B and Basal, where Luminal A and Luminal B signatures have 2 overlapping genes between them. The genes are up-regulated in Luminal B subtype and down-regulated in Luminal A. Since the signature-dependent crossover methods cannot inherently handle overlapping genes, we use an adaptation of these methods. To generate a new sample of type {Luminal A, Luminal B} using the crossover methods, only the label-defining signature includes the overlapping genes. To generate a Basal type that has no overlapping genes, one of the non-defining signatures is chosen at random to contribute the overlapping genes to maintain consistency in the expression pattern of the genes.

Figure 3 illustrates the distribution of balanced accuracy scores in the cross-validation test splits on the METABRIC external dataset. The Mod. Gamma-Poisson and Poisson methods show the best generalisation performance as the augmentation size increases. These methods are significantly better than all the other methods (except each other) and show an improvement of 8 − 9% over models trained with unaugmented data in classifying the subtypes. The adaptation of the crossover methods for handling overlapping genes does not work as evidenced by the poor generalisation performance. We hypothesize that the correlation between genes in the signatures of Luminal A and Luminal B prevent the crossover sampling methods from maintaining fidelity in the data. Detailed results on the METABRIC dataset is described in Appendix Table 13. Results on in-domain testing of the augmentation methods, including the original implementations of Gamma-Poisson and Poisson sampling, are illustrated in Appendix Figure 14 and Appendix Table 12 and lead to a similar conclusion. We also illustrate the distribution of the ROC-AUC scores across all splits in Appendix Figure 15 for in-domain and Appendix Figure 16 for out-of-domain test sets.

**Figure 3:**
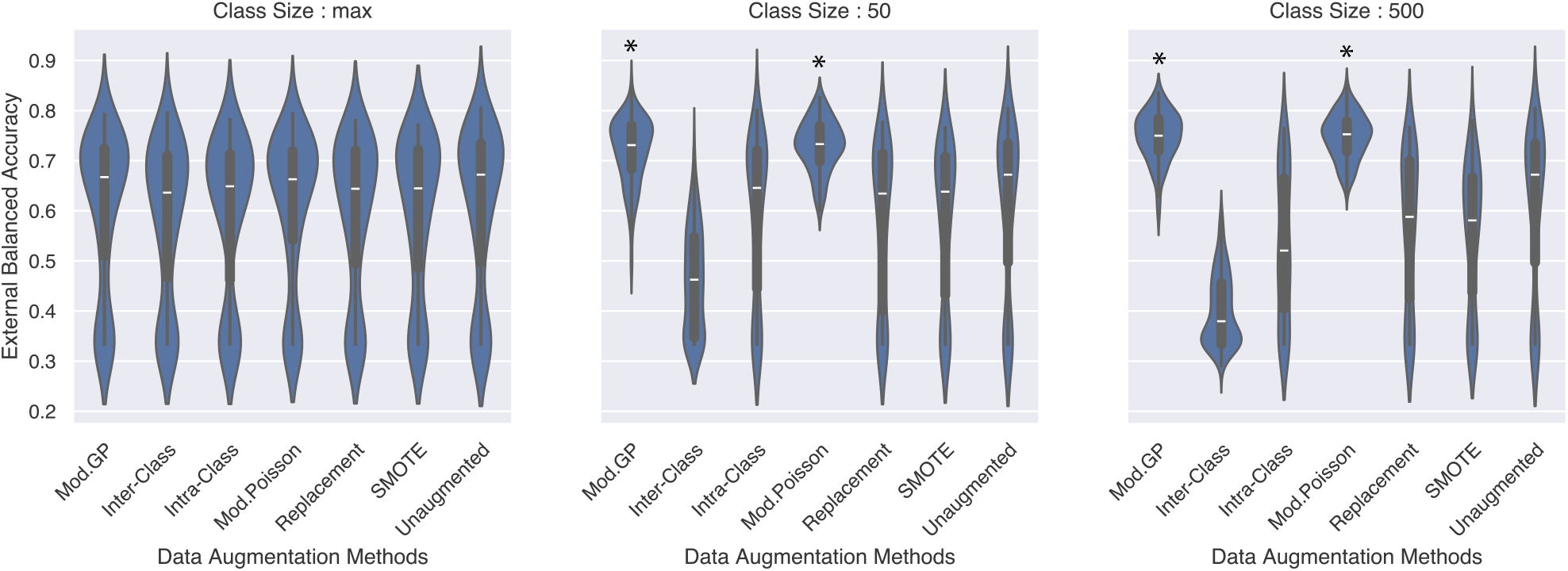
Distribution of balanced accuracy in the case of overlapping gene signatures on out-of-domain METABRIC dataset across all classification models (EBM, KNN, Logistic, RF, SVM-RBF) for every augmentation method, class size and CV split. The trend indicates that both Mod. Gamma-Poisson and Poisson methods achieve the best generalisation performance and are the best candidates when signatures overlap. * indicates that the methods are significantly better than all other methods except each other. We notice a bimodal distribution as SVM-RBF performs poorly, classifying samples at random (Appendix Table 13).

To summarise, the inter-class crossover method should be preferred when gene signatures do not overlap as evidenced by its significantly higher generalisation power (Figure 2). Should the gene signatures overlap, the Mod. Gamma-Poisson method is the better choice; its ability to capture over-dispersion in the data over Poisson-based sampling [14, 15] makes it preferable.

### 4.2 Effect of real data size

Figure 4 shows the in-domain averaged balanced accuracy across all classification models considered for the three class sizes at different percentages of real data in training. The trends in all three subplots are consistent with the fact that as we increase the number of real data observations, the performance of the classification models improve. We see again that over-augmenting the data only has a limited improvement for the distribution based methods. For instance, for the Mod. Gamma-Poisson method on 10% of the real training data, the associated average balanced accuracy increases from 82.444% ± 0.0528 to 82.54% ± 0.0571 when augmenting from class size 500 to 5000.

**Figure 4:**
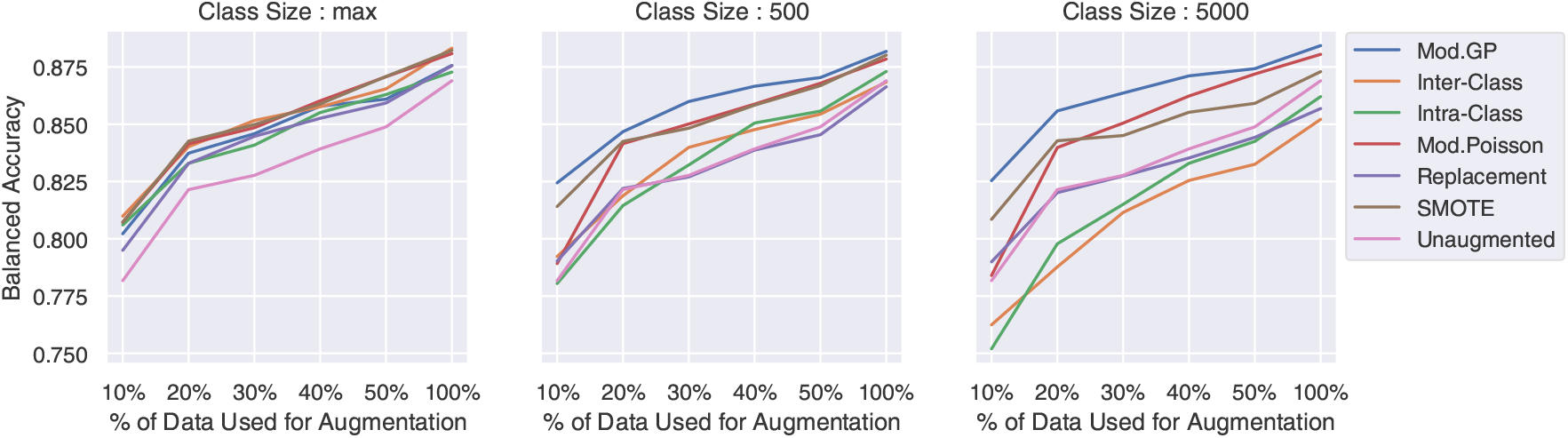
Trends in average classification performance on in-domain TCGA test set as the percentage of available real training data increases. Mod. Gamma-Poisson sampling is resistant to over-augmentation.

Figure 5 shows the out-of-domain averaged balanced accuracy across all classification models. At class sizes 500 and 5000, the inter-class crossover method achieves the best performance. At class size 500, this method achieves 65.55% ± 0.0492 when only 10% of the real training data is used, which is as high as that of using 100% of the data. The second best method is Mod. Gamma-Poisson, which achieves a similarly high performance but at class size 5000 (65.46% ± 0.0515).

**Figure 5:**
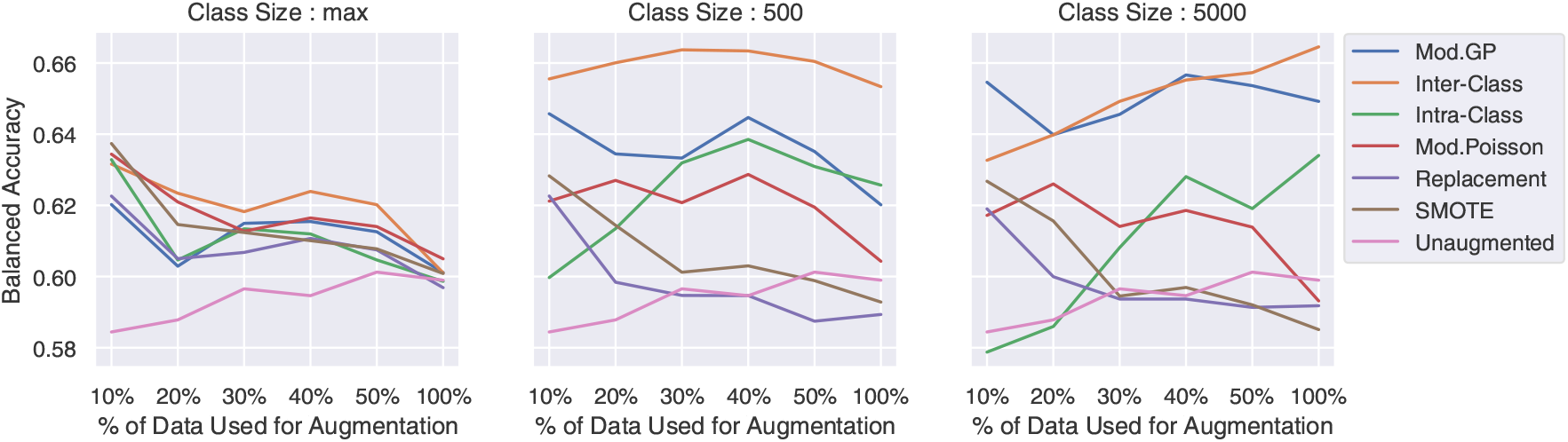
Trends in average classification performance on out-of-domain CPTAC dataset as the percentage of available real training data increases. Inter-class crossover sampling is resistant to over-augmentation and produces the best performance with fewer augmented data points compared to other methods.

In conclusion, the inter-class crossover method is the best for generalisation and over-augmentation does not significantly help performance other than adding to computational runtime, and, for some augmentation methods, it can even be detrimental to performance. We find that augmenting each class to size 500 proves to be optimal for all augmentation methods. Henceforth, we carry out further experiments using this class size.macro:

Figure 4 shows the in-domain averaged balanced accuracy across all classification models considered for the three class sizes at different percentages of real data in training. The trends in all three subplots are consistent with the fact that as we increase the number of real data observations, the performance of the classification models improve. We see again that over-augmenting the data only has a limited improvement for the distribution based methods. For instance, for the Mod. Gamma-Poisson method on 10% of the real training data, the associated average balanced accuracy increases from 82.444% ± 0.0528 to 82.54% ± 0.0571 when augmenting from class size 500 to 5000.

Figure 5 shows the out-of-domain averaged balanced accuracy across all classification models. At class sizes 500 and 5000, the inter-class crossover method achieves the best performance. At class size 500, this method achieves 65.55% ± 0.0492 when only 10% of the real training data is used, which is as high as that of using 100% of the data. The second best method is Mod. Gamma-Poisson, which achieves a similarly high performance but at class size 5000 (65.46% ± 0.0515).

In conclusion, the inter-class crossover method is the best for generalisation and over-augmentation does not significantly help performance other than adding to computational runtime, and, for some augmentation methods, it can even be detrimental to performance. We find that augmenting each class to size 500 proves to be optimal for all augmentation methods. Henceforth, we carry out further experiments using this class size.macro:

Using the colorectal cancer datasets, we evaluate the impact of initial dataset size on the different augmentation strategies. We select increasing percentages of the real training data - namely 10, 20, 30, 40 and 50 percent, in a stratified manner to reflect the class imbalances present in the original dataset. This simulates the “very low data regime” scenario. We repeated the classification experiment for each percentage of real data and tested the models on the in-domain TCGA COADREAD test set and unseen out-of-domain CPTAC COAD dataset at different augmented class sizes. Detailed results of individual classification models with each augmentation method and class size for the 10% real data scenario are shown in Appendix Table 14 and Table 15.

Figure 4 shows the in-domain averaged balanced accuracy across all classification models considered for the three class sizes at different percentages of real data in training. The trends in all three subplots are consistent with the fact that as we increase the number of real data observations, the performance of the classification models improve. We see again that over-augmenting the data only has a limited improvement for the distribution based methods. For instance, for the Mod. Gamma-Poisson method on 10% of the real training data, the associated average balanced accuracy increases from 82.444% ± 0.0528 to 82.54% ± 0.0571 when augmenting from class size 500 to 5000.

Figure 5 shows the out-of-domain averaged balanced accuracy across all classification models. At class sizes 500 and 5000, the inter-class crossover method achieves the best performance. At class size 500, this method achieves 65.55% ± 0.0492 when only 10% of the real training data is used, which is as high as that of using 100% of the data. The second best method is Mod. Gamma-Poisson, which achieves a similarly high performance but at class size 5000 (65.46% ± 0.0515).

In conclusion, the inter-class crossover method is the best for generalisation and over-augmentation does not significantly help performance other than adding to computational runtime, and, for some augmentation methods, it can even be detrimental to performance. We find that augmenting each class to size 500 proves to be optimal for all augmentation methods. Henceforth, we carry out further experiments using this class size.

### 4.3 Mixing gene blocks has added value

In this experiment, we analyse whether the crossover sampling strategy interferes with the biological signal of the sample. We consider the scenario where only 10% of the real training data (≈ 40 samples) is available. Since we retain the gene blocks as is, we do not destroy phenotype-specific information through mixing. We demonstrate this point by showing that key clinical variables of colorectal cancer subtypes, namely, microsatellite instability (MSI) and CpG Island Methylator Phenotype (CIMP), are still predictable. Predicting these variables, among others such as demographic factors, genomic markers and treatment response, is relevant for patient prognostics, disease monitoring and disease subtyping [35– 40]. Since we cannot infer MSI and CIMP labels for augmented data points, we first train a variational autoencoder using the augmented data (40-dimensional), retrieve the compressed embeddings (4-dimensional) of real data points and train classifiers on these embeddings. Thus, we indirectly evaluate the augmentation methods’ ability to preserve information from the subtypes.

Aggregated results are shown in Table 1, where we report the average balanced accuracy across all classifiers in predicting the MSI and CIMP status of patients. The inter-class and intra-class augmentation methods show a significant improvement over unaugmented data - 12% in-domain and 20% out-of-domain for MSI prediction, and 18% in-domain for CIMP prediction. These results prove that mixing of gene blocks between samples does not drastically affect the predictiveness of other variables associated with these colorectal cancer subtypes. Results for each classifier and augmentation method can be found in Appendix Table 18 and Table 19 for in-domain and out-of-domain MSI prediction, and Table 20 in-domain CIMP prediction.

**Table 1:**
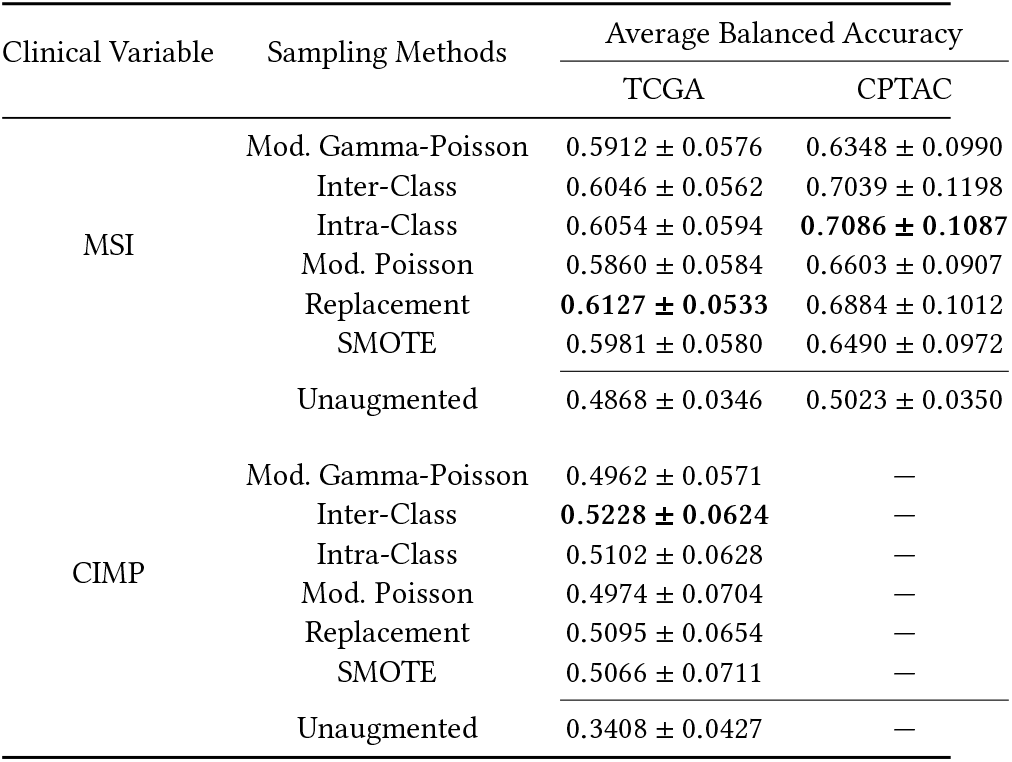
Average balanced accuracy scores and their standard deviation of a 5×5 cross-validation for MSI and CIMP Classification on class size 500 with only 10% real training data. The best scores are highlighted in bold. For both MSI and CIMP classification, our crossover methods are consistently in the top 2 in both in-domain (MSI and CIMP) and out-of-domain validation (MSI only). The results show that our crossover methods significantly improve prediction power over unaugmented data.

| Clinical Variable | Sampling Methods | Average Balanced Accuracy |  |
| --- | --- | --- | --- |
|  |  | TCGA | CPTAC |
| MSI | Mod. Gamma-Poisson | $0.5912 \pm 0.0576$ | $0.6348 \pm 0.0990$ |
| | Inter-Class | $0.6046 \pm 0.0562$ | $0.7039 \pm 0.1198$ |
| | Intra-Class | $0.6054 \pm 0.0594$ | **$0.7086 \pm 0.1087$** |
| | Mod. Poisson | $0.5860 \pm 0.0584$ | $0.6603 \pm 0.0907$ |
| | Replacement | **$0.6127 \pm 0.0533$** | $0.6884 \pm 0.1012$ |
| | SMOTE | $0.5981 \pm 0.0580$ | $0.6490 \pm 0.0972$ |
| | Unaugmented | $0.4868 \pm 0.0346$ | $0.5023 \pm 0.0350$ |
| CIMP | Mod. Gamma-Poisson | $0.4962 \pm 0.0571$ | — |
|  | Inter-Class | **$0.5228 \pm 0.0624$** | — |
| | Intra-Class | $0.5102 \pm 0.0628$ | — |
| | Mod. Poisson | $0.4974 \pm 0.0704$ | — |
| | Replacement | $0.5095 \pm 0.0654$ | — |
| | SMOTE | $0.5066 \pm 0.0711$ | — |
| | Unaugmented | $0.3408 \pm 0.0427$ | — |

**Table 2:**
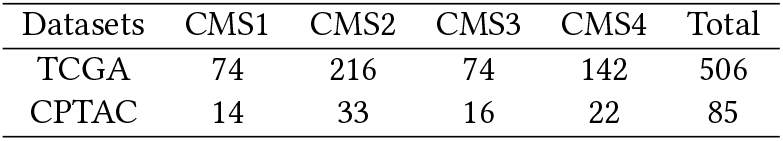
Sample counts of the different colorectal cancer sub-types in the TCGA and CPTAC gene expression datasets.

| Datasets | CMS1 | CMS2 | CMS3 | CMS4 | Total |
| --- | --- | --- | --- | --- | --- |
| TCGA | 74 | 216 | 74 | 142 | 506 |
| CPTAC | 14 | 33 | 16 | 22 | 85 |

### 4.4 Generation quality

Next, we evaluate the augmentation methods in the generative modeling task where only 10% of real training data is available. Here, we evaluate how effectively the augmentation contributes to estimating a parametric representation of the data. Our hypothesis is that greater diversity in the input samples will lead to more accurate parametric estimates of the underlying distribution. To verify this hypothesis, we extract test data embeddings from a pretrained variational autoencoder, setting class size to 500 and using 10% of the real training data. We compute the mean and standard deviation of these embeddings grouped by their true CMS class and initialise a Gaussian-Mixture model. From this model, we sample embeddings which are then decoded to generate new data. This procedure is done for both TCGA and CPTAC test data. These newly generated data are then passed through our previously trained classifiers from Section 4.1 studying generalisation performance.

A considerable performance difference is seen on the out-of-domain CPTAC dataset as shown in Figure 6. The inter-class crossover method (average balanced accuracy 66.36% ± 0.0631) performs the best over other methods, with an increase in accuracy of ≈ 35% over unaugmented data, ≈ 6% over Mod. Gamma-Poisson, Mod. Poisson and replacement methods, and ≈ 5% over SMOTE. The results of the experiment are shown in Appendix Table 16 and Table 17. A high classification accuracy indicates that the VAE was trained on data of sufficient quality to capture the underlying distribution. Appendix Figure 17 shows the average balanced accuracy achieved by each model on generated samples modeled on in-domain TCGA test set.

**Figure 6:**
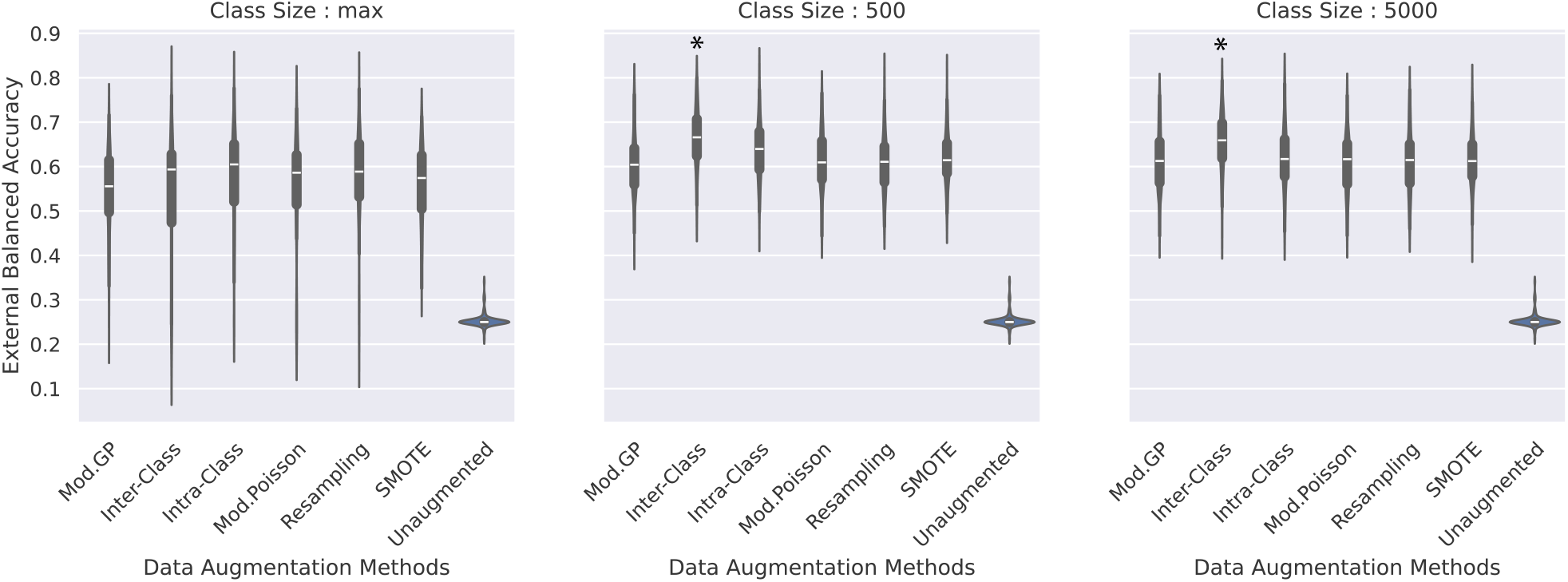
Classification performance on VAE generated samples modeled on out-of-domain CPTAC test set. On average, the inter-class crossover sampling is associated with the best performance (66.36% ± 0.0631). * indicates that the method is significantly better than all other methods.

### 4.5 Computational Complexity

In this section, we analyse the computational complexity of the augmentation methods. As the crossover methods sample signature blocks from a uniform distribution, both methods have a runtime complexity of 𝒪(*pk*) where *p* is the number of phenotypes and *k* is the number of new samples to be generated. Thus the complexity is linear in both k (for fixed p) and p (for fixed k). The parametric methods have a runtime complexity of 𝒪(*psnk*) + 𝒪(*k*), where *p* is the number of phenotypes, *s* = |*S*|, the number of genes in the signature, and *n* is the number of samples from which distribution parameters are computed. The first term is the complexity of computing distribution parameters, and the second term is the complexity of sampling values from the distribution. Since *s, p, n* and *k* are all independent from each other, the runtime complexity is linear. The random oversampling method has a runtime complexity of 𝒪(*kn*) since we generate *k* new samples and for each sample we go through the *n* values in the dataset for sampling. SMOTE has a quadratic runtime complexity since it requires the computation of a pairwise distance matrix, and is dependent on the number of samples, n, in the dataset (𝒪(*n*)^2^). Empirical runtimes are discussed in Appendix G.4.

## 5 Discussion

We introduce two classes of data augmentation strategies - signature-dependent and signature-independent methods. When distinct gene signatures represent target phenotypes, our signature-dependent crossover approach allows us to generate synthetic data that allows models to significantly generalise well to unseen, out-of-domain datasets. The reliance of these methods on distinct gene signatures is a limiting factor to its broad use. This could potentially be overcome by treating the gene signature blocks as dynamic instances that change size and content depending on the sample being generated. A deeper analysis into adaptations to handle overlapping genes could be a future research direction to make these methods versatile. Another potential limiting factor is the correlation of genes between two closely related phenotypes. Future research could investigate the effect of correlation on the sampling method. However, in the event that such signatures overlap, correlate or are unavailable entirely, our proposed modification of the Gamma-Poisson sampling method can be used. This signature-independent sampling method significantly outperforms other commonly used data augmentation methods in classification tasks. Although the Mod. Poisson sampling method sometimes performs on-par, particularly in the lower augmentation size regime, the Mod. Gamma-Poisson should be the preferred method as it can handle over-dispersion.

The findings related to effect of augmentation size on performance is an interesting byproduct of this study. From our experiments, we find that over-augmenting data beyond 1-2 orders of magnitude has little to no benefits. It typically leads to *<* 1 − 2% performance gain at increased computational cost, irrespective of the augmentation method. Our study shows that over-augmenting with generally well-performing methods like random oversampling and SMOTE can lead to worse generalisation performance, highlighting the importance of picking the right augmentation method on a case-basis.

Further research on extending the 2 classes of methods to the entire genome without phenotype conditioning would provide additional insights into their applicability to unsupervised learning. Our proposed sampling methods are intuitive, simple applications that are efficient and bring about better performance than state-of-the-art methods, holding promise for heterogeneity studies and disease analysis. By creating a landscape of mixed phenotypes, new opportunities would present itself in unsupervised and semi-supervised learning in scarce data regimes given fully unlabeled data.

## A Intra-class crossover sampling formalisation

Consider a dataset with n samples *x*^*i*^, where *i* ∈ 1, …, *n*, and m phenotype variants *P* ^*j*^, where *j* ∈ 1, …, *m*. Each sample *x*^*i*^ is of one phenotype *P* ^*j*^. Given each variant *P*^*j*^ is associated with a gene signature *S* ^*j*^, we represent each sample by the set of signatures associated with all possible variants, that is, *x*^*i*^ = {*S*^1^, …, *S*^*m*^}. The number of genes in a signature is given by |*S* ^*j*^ |. Therefore, each patient is represented by 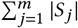 genes. For the sake of simplicity, the genes are ordered by signature - all genes belonging to signature *S*_1_ come first and so on, thus creating signature blocks, one for each phenotypic variant. To generate a new sample, *x*^*n* +1^ with phenotype *P* ^*j* =1^, we first subset all samples, belonging to this phenotype (Equation 3). This subset, *X* ^*j* =1^, is analogous to a bag of signature blocks (Equation 4).

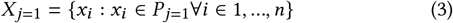

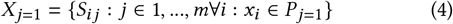

We randomly sample signature blocks from this subset (Equation 5) which are put together to create a new sample *x*^*n*+1^.

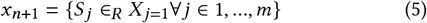

We illustrate a practical example on the gene expression of patients with colorectal cancer, using the Consensus Molecular Subtype (CMS) as phenotype in subsubsection 2.1.1.

## B Inter-class crossover sampling formalisation

Consider a similar premise as the one described in subsubsection 2.1.1. To generate a synthetic sample of phenotype *P* ^*j* =1^, we sample *S* ^*j* =1^ only from the subset of patients, *X* ^*j* =1^, that have phenotype *P* ^*j* =1^, that is. *S* ^*j*^ ∈^*R*^ *X* ^*j*^. We then randomly sample the remainder signatures *S*^*k*^ for *k* ∈ 1, …, *m* and *k* ≠ *j*, from all samples except those that belong to phenotype *P*^*k*^ as described in Equation 6. The sampled signatures are then put together to create a new synthetic sample, *x*^*n*+1^ = {*S* ^*j*^, *S*^*k*^} for *k* ∈ 1, …, *m* and *k* ≠ *j* and where *j* = 1.

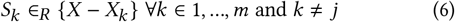

## C Modified Gamma-Poisson sampling

The negative-binomial (NB) distribution is a commonly used parametric method for augmenting RNA-Seq data for differential expression analysis due to it’s ability to capture the overdispersion typically seen in these data. To model the RNA-Seq read frequency *Y* of a given gene *j*, the distribution relies on two parameters, namely the mean and dispersion of the counts, i.e, *Y*^*j*^ = NB (*μ* ^*j*^, *ϕ*).

Here, *ϕ* is the dispersion parameter that controls the variance of the counts, 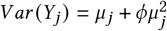. The estimation of *ϕ* is, however, not trivial. The negative-binomial (NB) distribution is typically expressed as a Gamma-Poisson mixture [25, 26] to make it more tractable. In this formulation, the rate parameter *λ* of the Poisson distribution is a random variable sampled from a gamma distribution parameterised by the shape (*α*) and rate (*β*) parameters, which are estimated from the observed population, *λ* ∼ Γ (*α, β*). The RNA-Seq read counts are then sampled from this initialised Poisson distribution, *Y*^*j*^ ∼ *Poisson* (*λ*^*j*^).

To ensure that the newly generated samples add some diversity while maintaining the same distribution as the augmented class, we create a mixture of Gamma-Poisson distributions. In this strategy, to generate every new observation, a subset *S* of samples is randomly chosen to estimate the *α* and *β* parameters from its mean *μ* and variance *σ*^2^ as shown in Equation 1. These parameters initialise the gamma distribution. Since the augmentation is aimed at solving class imbalance, the subset is created from samples belonging to the same class. Thus, the newly generated observation will have the same label as the subset. The size of the subset |*S*| is a hyper-parameter defined by the user. The random variable sampled from this gamma distribution is used to initialise the Poisson distribution from which a new observation is sampled. This process is repeated *n* times to generate *n* new observations. We recommend a smaller subset size to maximise the variance in the subset. A larger variance would mean more scattered observations, thereby generating a diverse set of samples as opposed to dense local clusters with a smaller variance.

## D Modified Poisson sampling

The Poisson distribution is another method commonly used to model RNA-Seq read counts. Similarly to the Gamma-Poisson strategy, a subset is first sampled from the class to be augmented, after which the mean is computed and set as the rate parameter *λ* for the Poisson distribution as shown in Equation 2.

However, unlike the Gamma-Poisson, this distribution assumes the variance of the sample equals the mean and fails to capture overdispersion in the data. To generate new observations with this method, a larger subset size is recommended such that the variance is closer to the mean while still being high enough to generate variability in the new data.

## E Visualising augmented data for other class sizes

Figure 9 illustrates the UMAP projected distributions of the datasets augmented by various methods to class size 5000. These visualisations are compared to the UMAP projection of the unaugmented dataset to ascertain visually if global and local trends are fairly captured. Similar visualisation are also show for class size Max in Figure 7 and class size 500 in Figure 8.

**Figure 7:**
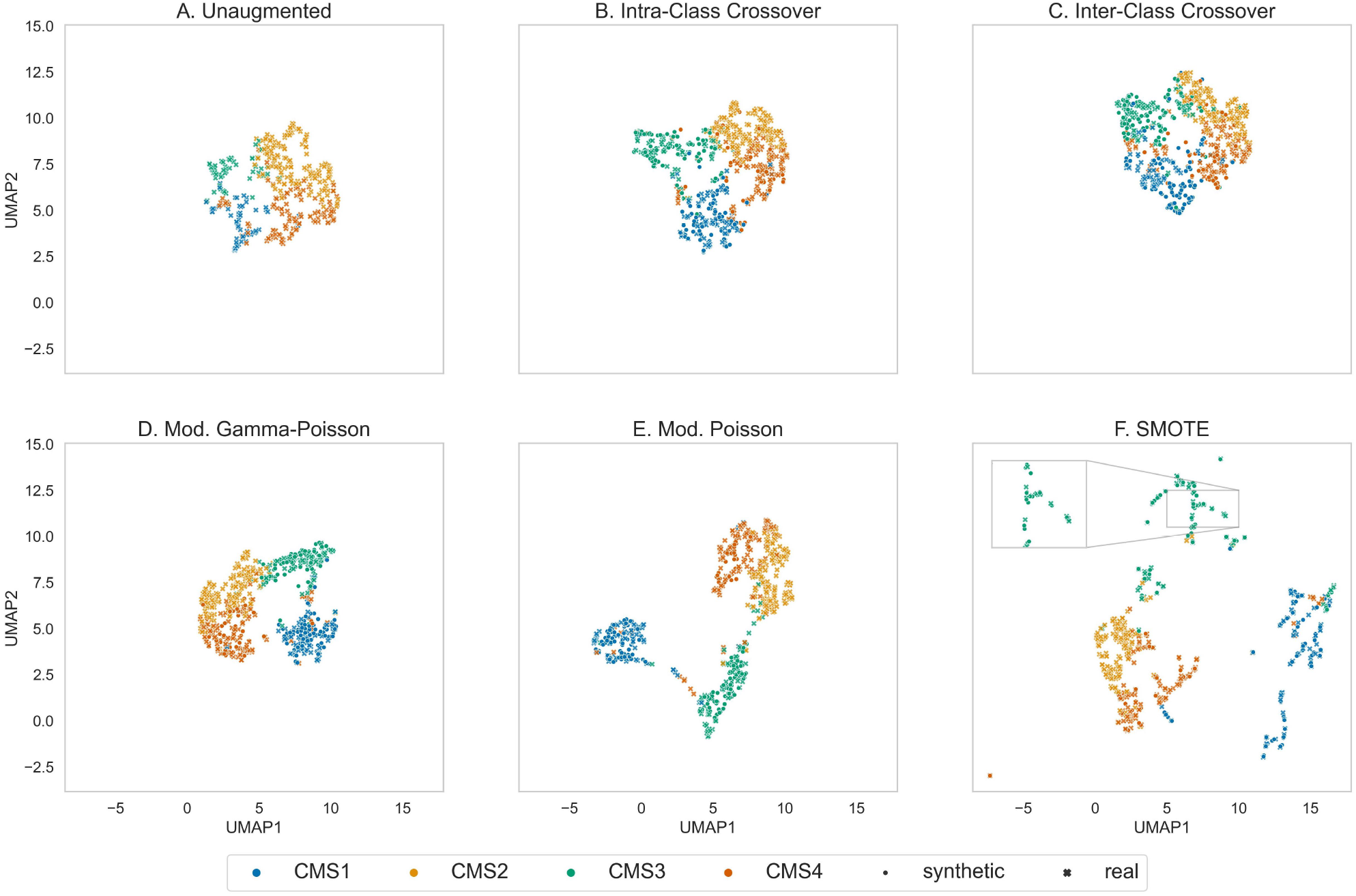
UMAP [41] visualisations of datasets augmented to match the majority class size. The intra-class and inter-class crossover methods capture both the global and local trends in the original unaugmented data better than the other methods. Mod. Gamma-Poisson and SMOTE closely follow. The Mod. Poisson sampling method begins to separate clusters which become more apparent and amplified as the class augmentation size is increased.

**Figure 8:**
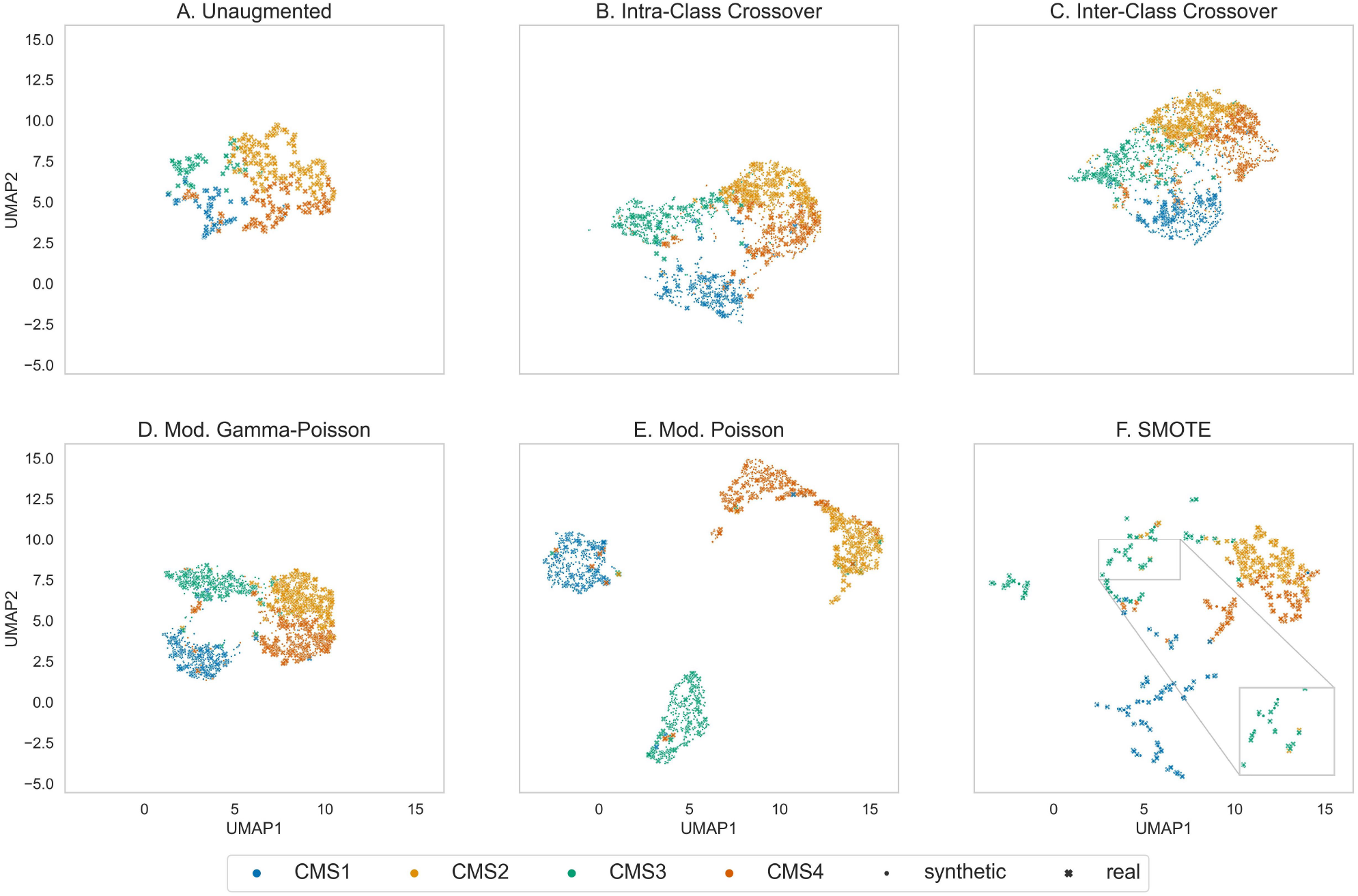
UMAP [41] visualisations of datasets augmented to have 500 samples per class. Data augmented by the intra-class, inter-class crossover methods and Mod. Gamma-Poisson sampling maintain a structure similar to the original unaugmented data. Data generated by the Mod. Poisson sampling method begins to show cluster separation, which could lead to spurious results. SMOTE generates samples on the line between two reference points and thus cannot populate sub-regions densely.

**Figure 9:**
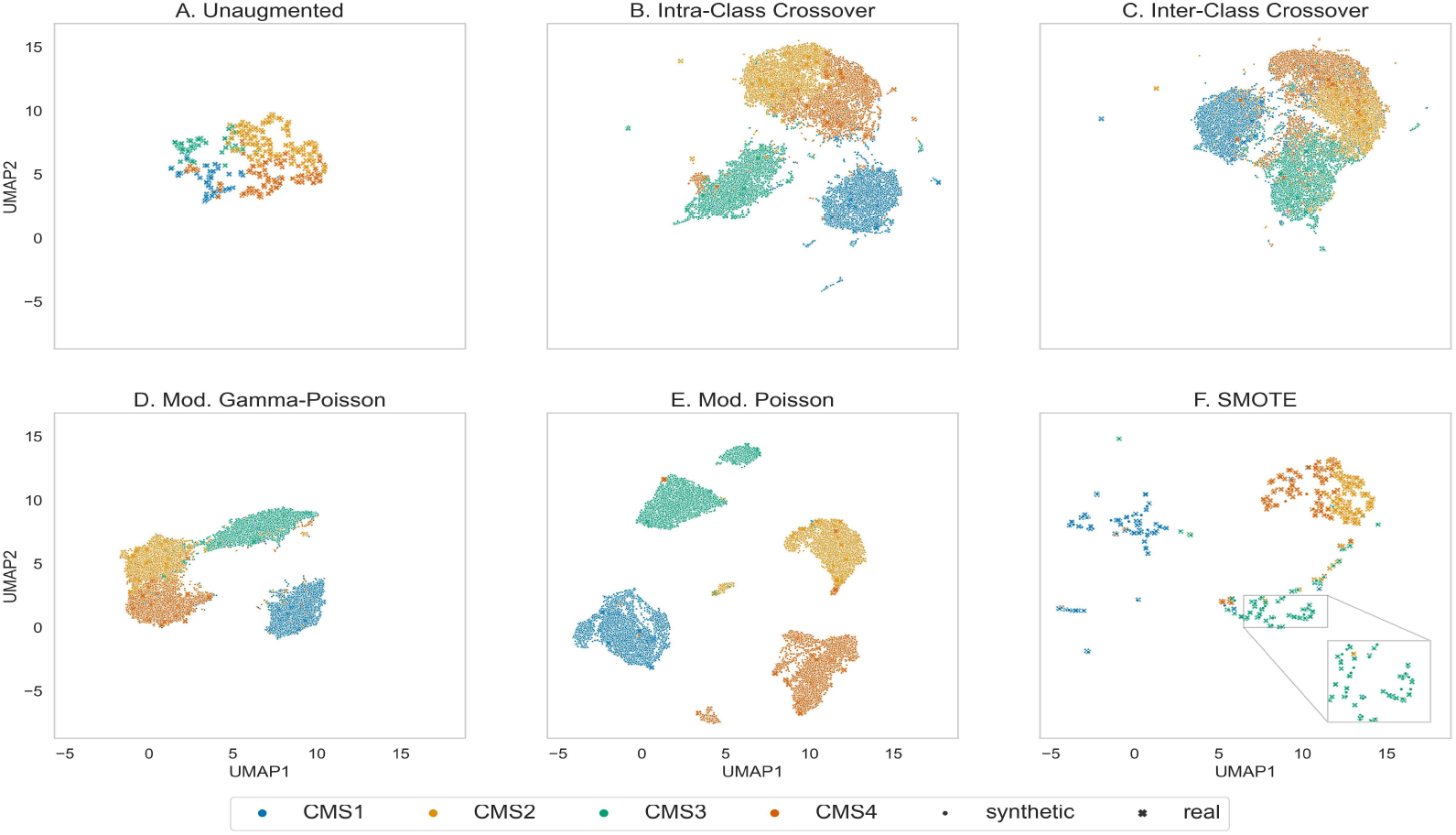
UMAP [41] visualisations of the augmented datasets show that the inter-class crossover sampling method best captures both the global and local structure of the original unaugmented data. All augmented datasets have 5000 samples per class.

## F Distribution size effect

In this experiment, we test various reference set sizes to understand its impact on CMS classification performance. The reference set sizes considered are *r* = 3, 5, 10, 25, 50. We choose *class*_*size* = 5000 so that there are enough newly sampled observations to draw a conclusion about the effect of reference set size. The study is conducted on a classification task with various classifiers and the 5-fold cross validation balanced accuracy is reported in Table 3. The reference set size with the highest accuracy on average, *r* = 5, is chosen for the Gamma-Poisson sampling method. Figure 10 illustrates what the distribution looks like with different reference set sizes. We observe that for *r* = 5, the real and synthetic samples are well interspersed with each other showing qualitatively that the distribution of synthetic samples is similar to that of the real samples. As *r* increases, new samples are created that seemingly fall away from its member class resulting in new sub-clusters as seen in Figure 10 c and d, for CMS1 and CMS2 classes.

**Table 3:**
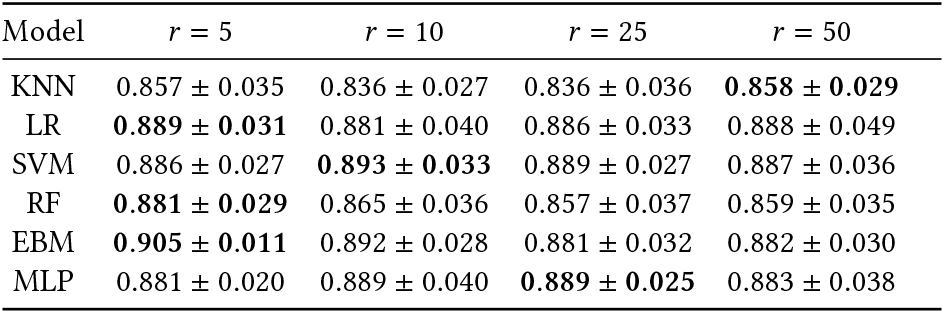
Effect of reference sample size on classifier performance for parametric methods. The table shows 5-fold cross-validation results including standard deviation for different *r* sizes and classifiers. Majority of the best performances belong to *r* = 5, which also has the best performance on average.

**Figure 10:**
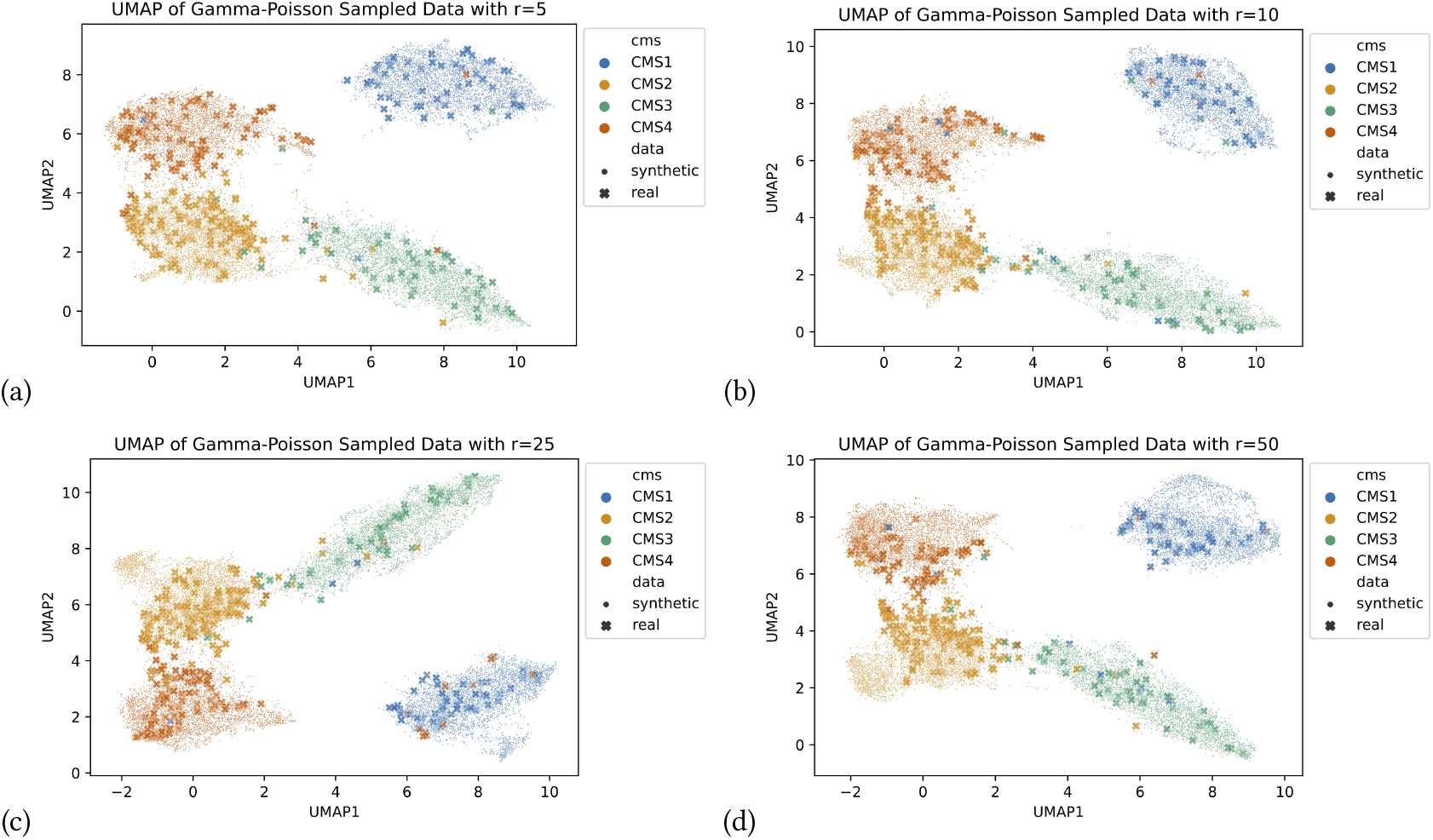
UMAP [41] visualisations of Gamma-Poisson augmented datasets in the study. (a)r=5. (b) r=10. (c) r=25. (d) r=50. All augmented datasets have 5000 samples per class. When r is set to 5, the new samples are more interspersed with the real data, as opposed to r=50, where new samples are generated “out of distribution” resulting in new clusters.

For the Poisson distribution, we double the reference set size to *r* = 10 such that the variance is more stable and closer to the mean. Figure 11 illustrates the Poisson augmented data at different reference set sizes. Here, we make a similar observation that as *r* increases, sub-clusters begin to form due to synthetic samples being generated well outside its member class distribution. On comparing the Poisson augmented data to the Gamma-Poisson augmented data, we observe that the CMS clusters appear to be well separated with the Poisson. While this may appear to be beneficial for certain tasks that are strictly associated with the these types, it does not capture the distribution of the real data well (Figure 9a). This could introduce spurious associations and attributions.

**Figure 11:**
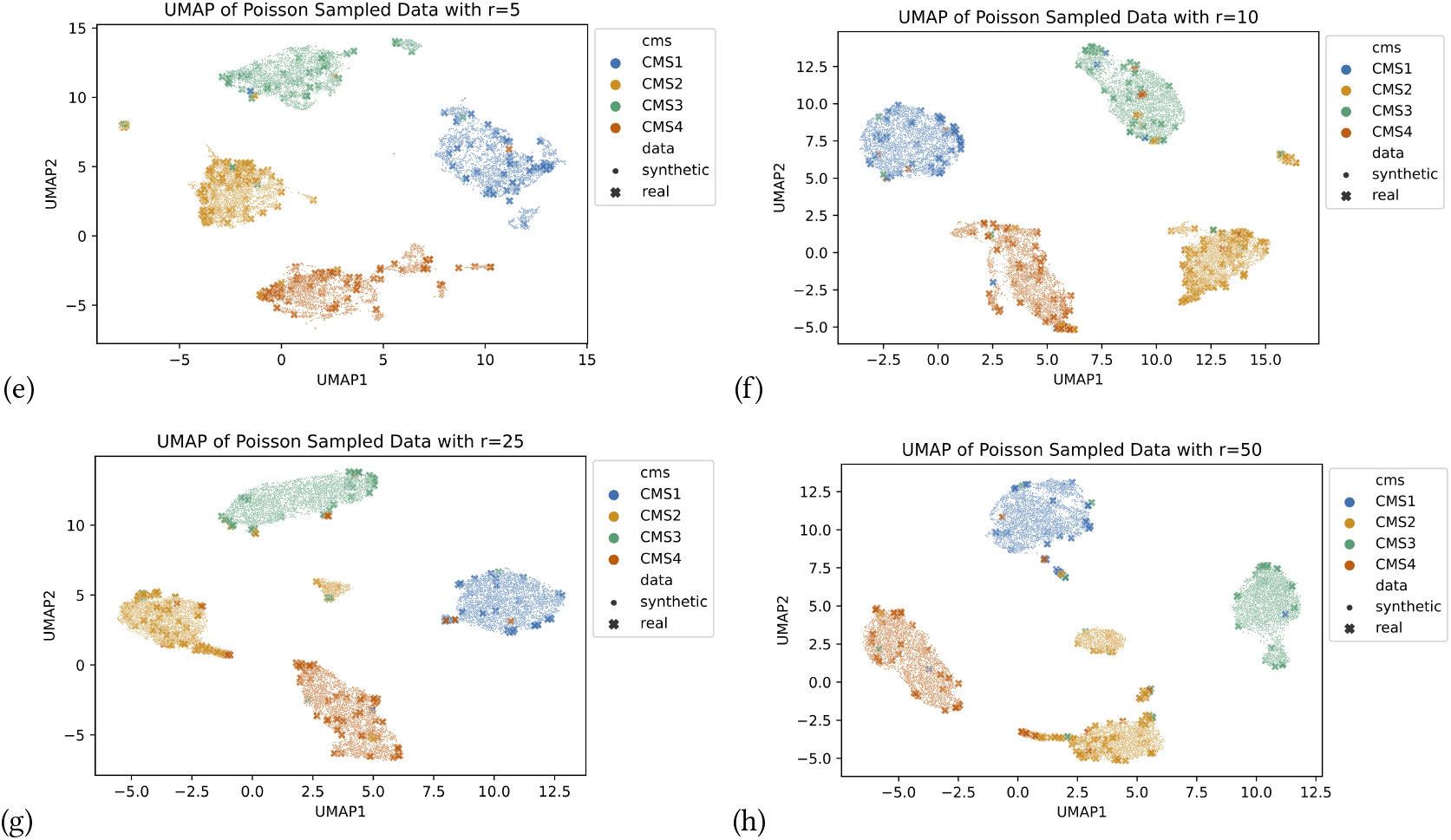
UMAP [41] visualisations of Poisson augmented datasets in the study. (a)r=5. (b) r=10. (c) r=25. (d) r=50. All augmented datasets have 5000 samples per class. For r=10, the new samples appear better interspersed with the real data and there are fewer outlier clusters.

**Figure 12:**
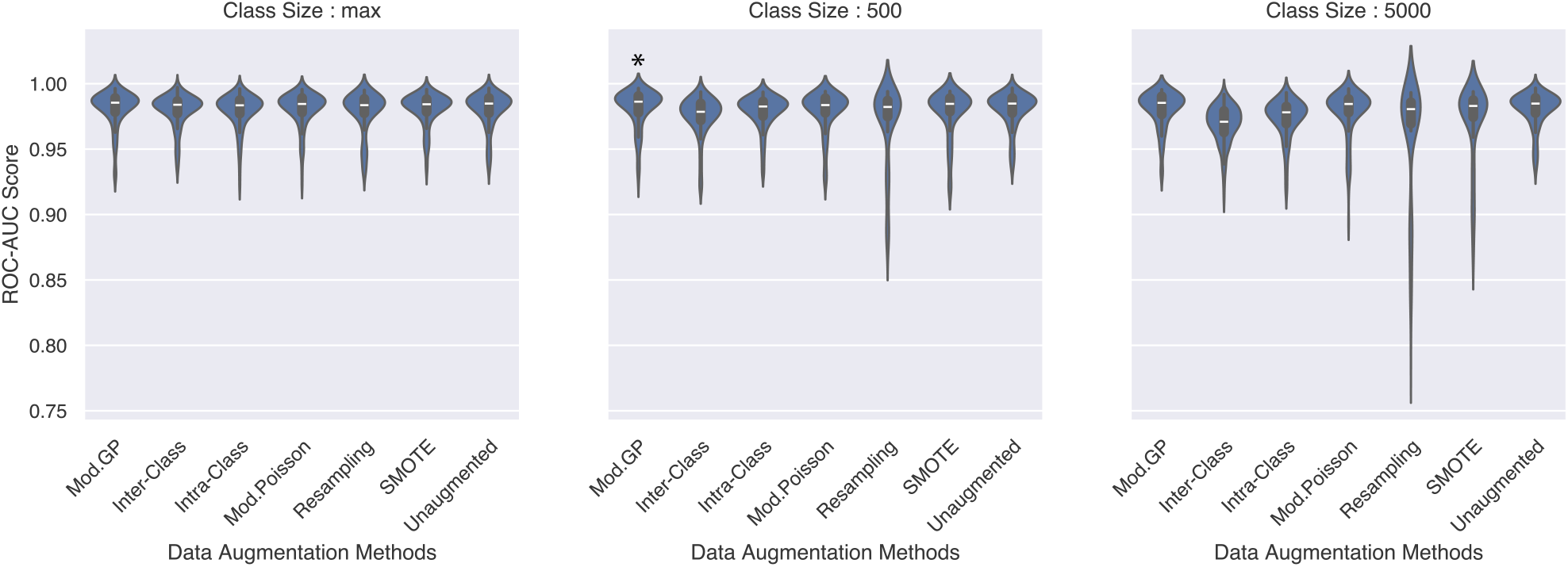
Distribution of ROC-AUC scores on in-domain TCGA test set across all classification models (EBM, KNN, Logistic, RF, SVM-RBF) for each augmentation method and class size. The trend indicates that as augmentation size increases, the Mod. Gamma-Poisson tends to achieve a higher score, while methods like SMOTE and replacement sampling perform worse. * indicates that the method is significantly better than the rest.

**Figure 13:**
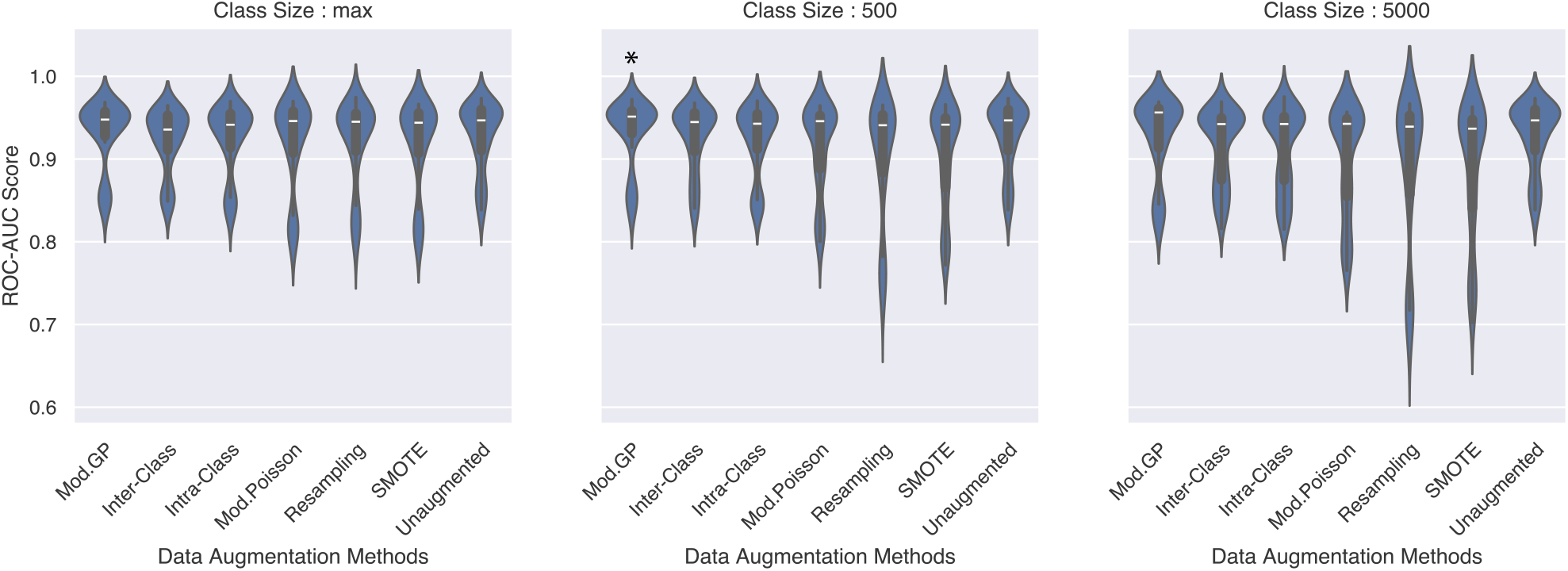
Distribution of ROC-AUC scores on out-of-domain CPTAC test set across all classification models (EBM, KNN, Logistic, RF, SVM-RBF) for each augmentation method and class size. * indicates that the method is significantly better than all the other methods. Replacement, SMOTE and Mod. Poisson are significantly worse than the other methods, including unaugmented, in all class sizes.

**Figure 14:**
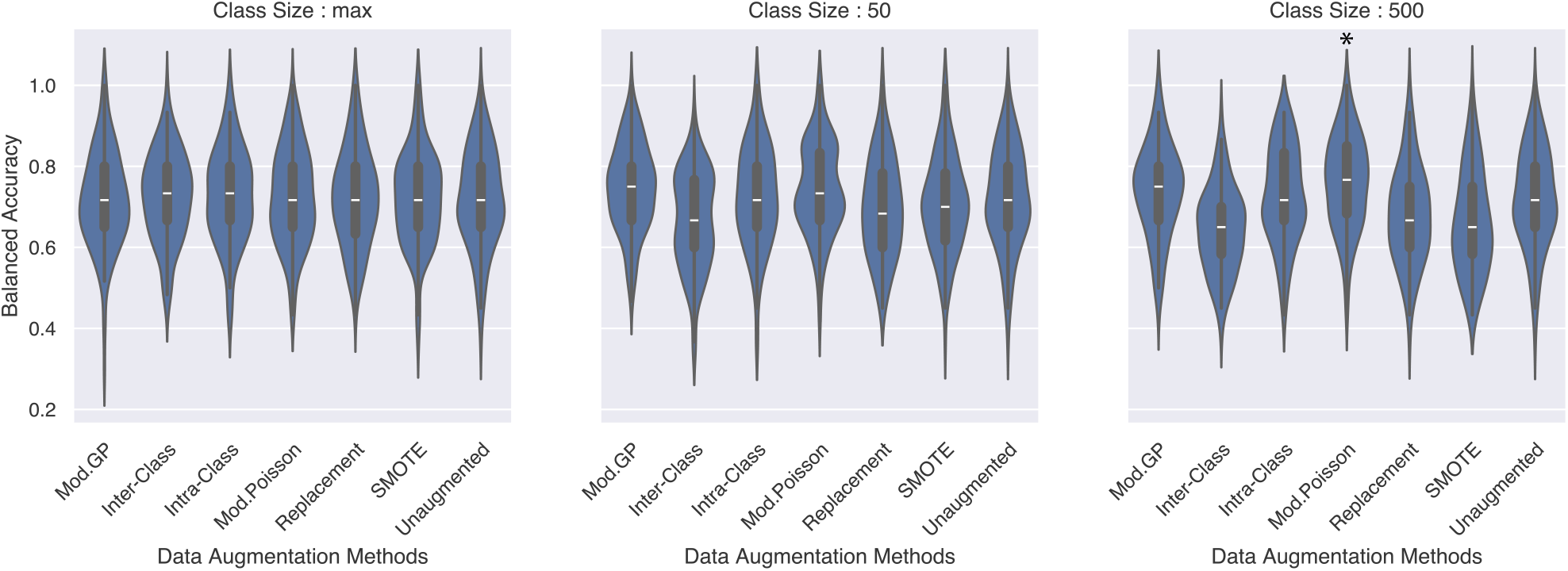
Distribution of balanced accuracy in the case of overlapping gene signatures on in-domain GSE20713 dataset across all classification models (EBM, KNN, Logistic, RF, SVM-RBF) for every augmentation method, class size and CV split. * indicates that the methods are significantly better than all other methods.

**Figure 15:**
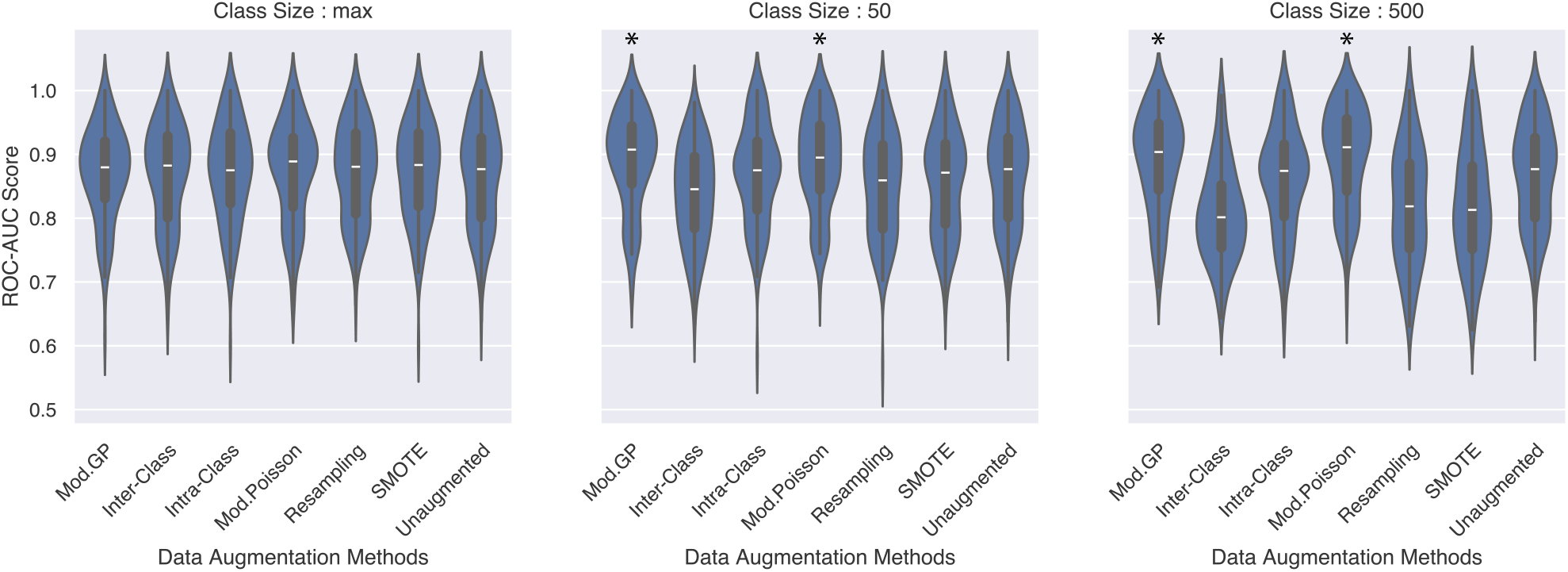
Distribution of ROC-AUC scores on in-domain GSE20713 test set across all classification models (EBM, KNN, Logistic, RF, SVM-RBF) for each augmentation method and class size. The trend indicates that as augmentation size increases, the Mod. Gamma-Poisson and Mod. Poisson methods tend to achieve a higher score, while methods like inter-class crossover, SMOTE and replacement sampling perform worse. * indicates that the method is significantly better than the rest, except each other.

**Figure 16:**
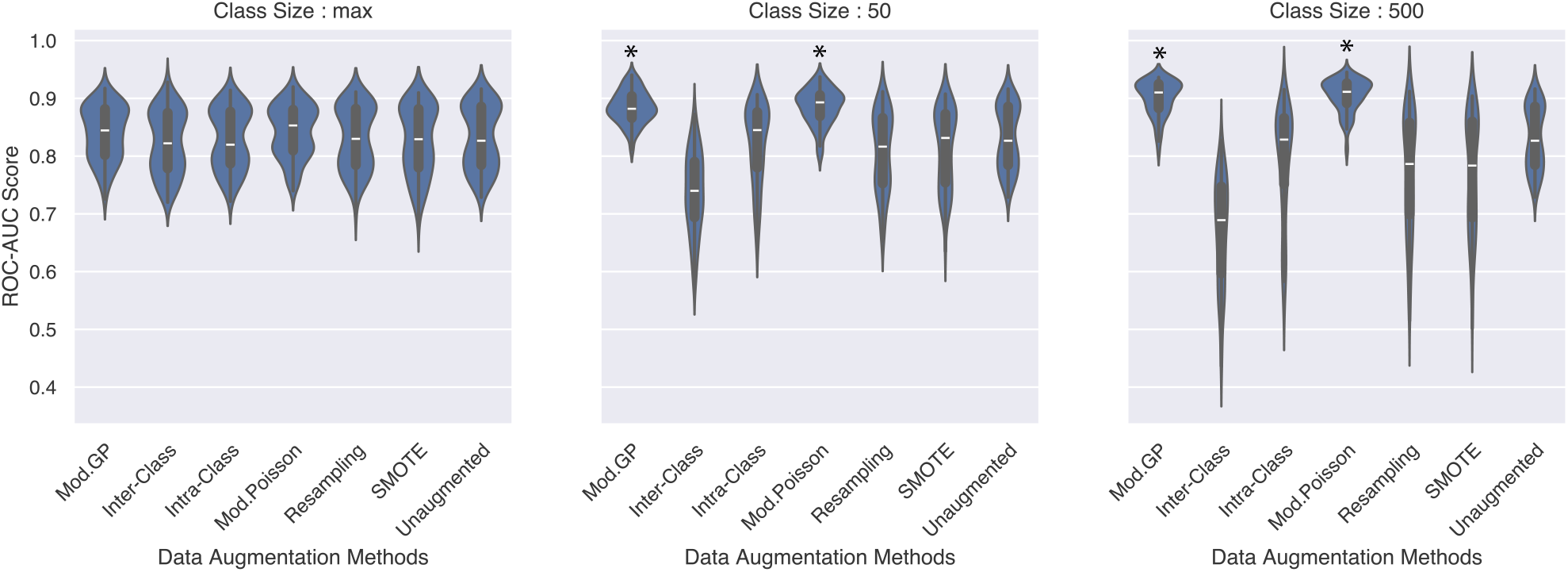
Distribution of ROC-AUC scores on out-of-domain METABRIC test set across all classification models (EBM, KNN, Logistic, RF, SVM-RBF) for each augmentation method and class size. The trend indicates that as augmentation size increases, the Mod. Gamma-Poisson and Mod. Poisson methods tend to achieve a higher score, while methods like inter-class crossover, SMOTE and replacement sampling perform worse. * indicates that the method is significantly better than the rest, except each other.

**Figure 17:**
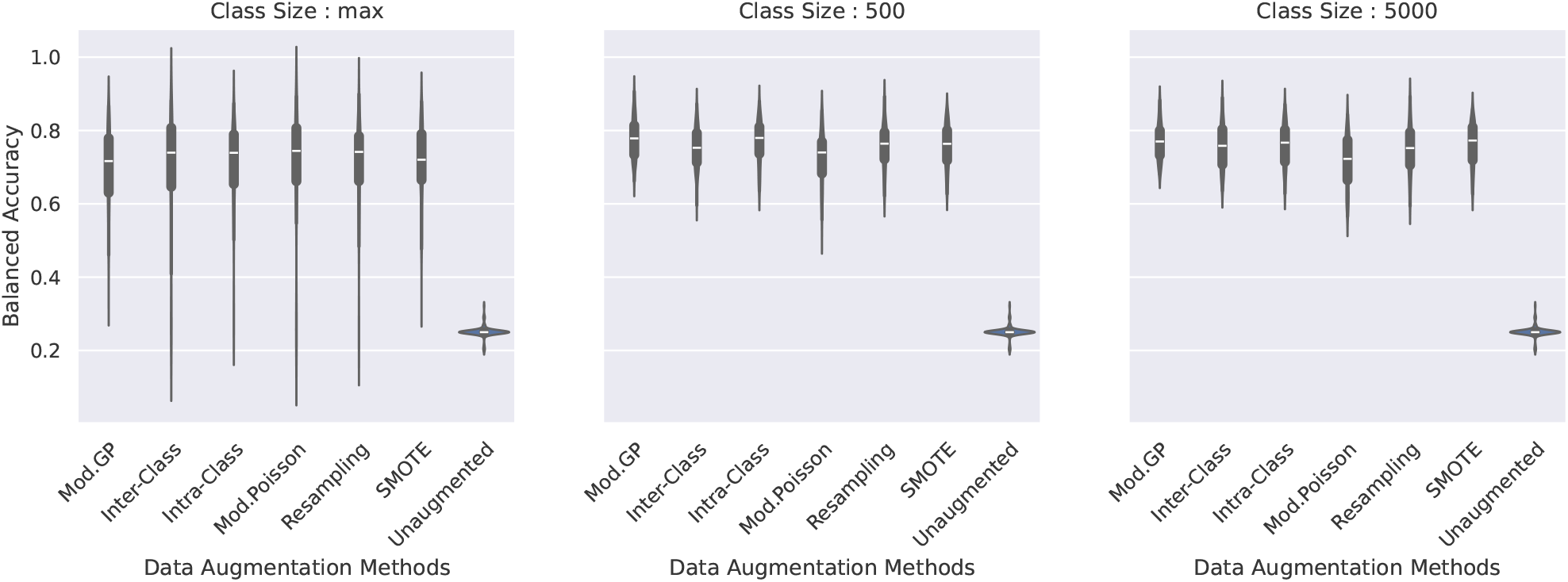
Classification performance on VAE generated samples modeled on in-domain TCGA test set. Although the methods show a similar performance, on average, the Mod. Gamma-Poisson augmentation method is associated with the best score(77.71% ± 0.0536). The VAEs generating the samples were trained on data augmented by the different methods as shown in the figure.

While we acknowledge that conclusions drawn from UMAP visualisations have to be taken with a grain of salt, we note that all UMAPs were generated with the same hyperparameter choice, differing only in their input data. This gives us a qualitative insight into what these differently augmented datasets with various reference set sizes look like and whether or not the newly generated data improve the support of a given class.

We omit results for *r* = 3 since it has high failure rates. Since the parametric distributions depend on the mean and variance of the subsample, it can sometimes happen that when *r* = 3 the variance associated with the subsample is too small, resulting in NaNs being sampled from the gamma distribution due to an invalid argument (negative integers or 0 possibly due to a possible overflow error).

## G Implementation details

### G.1 Classification performance

To demonstrate the utility of our augmentation methods in discriminative modelling, we make use of 5 classifiers, namely, Logistic Regression (LR), K-Nearest Neighbours (KNN), Support Vector Machines (SVM), Explainable Boosting Machines (EBM), and Random Forests (RF). For the standard classifiers, we make use of the `scikit-learn` [42] library implementations with the default parameters. We only increase the max_iter to 5000 for the Logistic Regression model. For the EBMs, we use the `InterpretML` library. We set a seed (42) for all models.

We report the mean balanced accuracy (Equation 7) on the TCGA standardised test sets and the CPTAC dataset. The metric is computed using true positive (TP), true negative (TN), false positive (FP) and false negative (FN) counts and is averaged across the 25 splits (5-fold CV repeated 5 times). The CPTAC dataset is standardised according to the training data mean and standard deviation from each split after which it is passed to the trained model from that split. We drop patients for whom the target label is not available. Additionally, we perform significance testing with the Wilcoxon test. All experiments are run locally on a 2019 MacBook Pro.

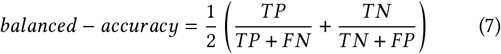

#### G.1.1 Effect of Data Size

The above process is repeated for augmented data containing various percentages of real data from 10 to 50 percent and 100 percent.

#### G.1.1 Predicting Other Variables

For the classification tasks of predicting MSI and CIMP status, we use the SVM-RBF and EBM models from `scikit-learn` and `InterpretML` to run our experiments (with the default parameters and seed 42). Here the input to the models is a 4D latent representation extracted from the VAE (described in subsection G.3) for each patient. Similar to the survival analysis task, we perform training and testing with the same 5×5 stratified cross-validation splits, and all samples for which the MSI/CIMP status is not available are dropped for the MSI/CIMP prediction task respectively. The patient summary counts for both MSI and CIMP are reported in Table 4.

**Table 4:** Summary counts of patients with MSI and CIMP labels available in the TCGA COADREAD and CPTAC COAD gene expression datasets.

| Clinical Variable | Categories | TCGA | CPTAC |
| --- | --- | --- | --- |
| MSI | MSI-H | 80 | 24 |
|  | MSI-L | 74 | - |
|  | MSS | 339 | 81 |
|  | Total | 493 | 105 |
| CIMP | CIMP.High | 66 | - |
|  | CIMP.Low | 86 | - |
|  | CIMP.Neg | 272 | - |
|  | Total | 364 | - |

**Table 5:** Sample counts of the different breast cancer subtypes in the TCGA and METABRIC gene expression datasets.

| Datasets | LuminalA | LuminalB | Basal | Total |
| --- | --- | --- | --- | --- |
| GSE20713 | 23 | 22 | 22 | 67 |
| METABRIC | 700 | 475 | 209 | 1384 |

**Table 6:** Empirical runtimes of the different augmentation methods averaged over 25 splits.

| Sampling Method | Time (seconds) |
| --- | --- |
| Mod. Gamma-Poisson | $136.348 \pm 2.321$ |
| Inter-Crossover | $0.642 \pm 0.028$ |
| Intra-Crossover | $0.066 \pm 0.004$ |
| Mod. Poisson | $60.874 \pm 0.824$ |
| Replacement | $0.029 \pm 0.003$ |
| SMOTE | $0.151 \pm 0.049$ |

### G.2 Handling overlapping genes

The sample counts for GSE20713 and METABRIC breast cancer datasets are described in Table 5.

We use GSE20713 as the training dataset, despite its low sample number, because it presents the perfect scenario of a dataset lacking in samples, thereby warranting augmentation.

To demonstrate the utility of our augmentation methods in handling overlapping gene signatures, we design a classification experiment using the methods described in section 3.3, namely, Logistic Regression (LR), K-Nearest Neighbours (KNN), Support Vector Machines (SVM), Explainable Boosting Machines (EBM), and Random Forests (RF). For the standard classifiers, we make use of the `scikit-learn` [42] library implementations with the default parameters. We only increase the `max_iter` to 5000 for the Logistic Regression model. For the EBMs, we use the `InterpretML` library. We set a seed (42) for all models.

We report the mean balanced accuracy (Equation 7) on the GSE20713 standardised test sets and the METABRIC microarray dataset. The metric is computed using true positive (TP), true negative (TN), false positive (FP) and false negative (FN) counts and is averaged across the 25 splits (5-fold CV repeated 5 times). The METABRIC dataset is standardised according to the GSE20713 training data mean and standard deviation from each split after which it is passed to the trained model from that split. We drop patients for whom the target label is not available. Additionally, we perform significance testing with the Wilcoxon test. All experiments are run locally on a 2019 MacBook Pro.

### G.3 Generation quality

The encoder and decoder of the VAE are single-layer neural networks with 20 hidden units activated by the ReLU function [43]. The dimension of the latent representation is 4. The batch size and learning rate are adapted according to the size of the dataset. For the unaugmented dataset and for the augmented datasets with class sizes ‘Max’ and ‘500’, we set the batch size to 64 and learning rate to 0.001. For all the other datasets, we set the batch size to 128 and learning rate to 0.0001. The models are trained on the sum of the L2 loss and KL-divergence. An L2 penalty of 0.0001 is applied to all models, which are optimised by Adam [44] and trained for 300 epochs with an early stopping wait time of 8 steps. The experiments to train a VAE were run on CPUs with 24 cores. The experiment for generating new samples and classifying them were run on 12 core CPUs.

### G.4 Empirical runtime

In this experiment, we measured the time taken to generate an augmented colorectal cancer dataset of class size 5000 for each of the 4 CMS classes given 100% of the original data (≈ 450 samples). The runtime is measured for each of the 25 cross validation splits and averaged. We would like to note that our code implementations of the signature-dependent and signature-independent methods may not be efficient and so these methods (except intra-class crossover) appear to be slower than SMOTE.

#### G.4.1 Scalability of crossover methods

Empirical results show that a 10-fold increase in dataset size results in a 2.69% increase in time taken to generate the dataset (Table 7). We measured the time taken to generate an augmented colorectal cancer dataset of class size 5000 for each of the 4 CMS classes in 2 scenarios (repeated 3 times), namely, given 10% of the original data (≈ 40 samples) and given 100% of the original data (≈ 450 samples). These results suggest that our method is scalable to larger datasets without incurring drastic computational costs. Optimising the code could achiever faster runtimes.

**Table 7:** Average computational time for the crossover techniques to generate a colorectal cancer dataset of class size 5000 for each of the 4 CMS classes (repeated 3 times) when given 10% (≈ 40 samples) and 100% of the data (≈ 450 samples).

| Crossover Type | Computational Time (seconds) |  |
| --- | --- | --- |
|  | 10% of Data | 100% of Data |
| Inter-Class Crossover | 0.6039 | 0.6202 |
| Intra-Class Crossover | 0.0707 | 0.0803 |

## H Extended results

In this section, we include long-form tables that quantify metrics for figures in the main paper as well as supporting and additional results for the experiments conducted. The aggregated metrics are reported along with their standard deviation.

### H.1 Unmodified GP and Poisson methods perform worse

Table 8 describes the results of CMS classification of the unmodified GP and Poisson methods on in-domain TCGA test set. Table 9 describes the results of CMS classification of the unmodified GP and Poisson methods on out-of-domain CPTAC test set. Comparing results from Table 8 and Table 9 to Table 10 and Table 11, respectively, we see how the unmodified version of Gamma-Poisson and Poisson show poorer performance compared to the unmodified versions.

**Table 8:** Average 5×5 cross validation balanced accuracy and standard deviation on in-domain TCGA test set for the unmodified Gamma-Poisson and Poisson augmentation methods. The scores are reported for different classifier models and different class sizes. The data was augmented given 10% of the original dataset.

| Class Size | Sampling Methods | EBM | KNN | Logistic | RF | SVM-RBF | Average |
| --- | --- | --- | --- | --- | --- | --- | --- |
| max | Gamma-Poisson | 0.7913± 0.0519 | 0.7913± 0.0519 | 0.7913± 0.0519 | 0.7913± 0.0519 | 0.7913± 0.0519 | 0.7913± 0.0519 |
|  | Poisson | 0.8009± 0.0563 | 0.8009± 0.0563 | 0.8009± 0.0563 | 0.8009± 0.0563 | 0.8009± 0.0563 | 0.8009± 0.0563 |
| 500 | Gamma-Poisson | 0.7982± 0.0460 | 0.7982± 0.0460 | 0.7982± 0.0460 | 0.7982± 0.0460 | 0.7982± 0.0460 | 0.7982± 0.0460 |
|  | Poisson | 0.8196± 0.0615 | 0.8196± 0.0615 | 0.8196± 0.0615 | 0.8196± 0.0615 | 0.8196± 0.0615 | 0.8196± 0.0615 |
| 5000 | Gamma-Poisson | 0.7972± 0.0574 | 0.7972± 0.0574 | 0.7972± 0.0574 | 0.7972± 0.0574 | 0.7972± 0.0574 | 0.7972± 0.0574 |
|  | Poisson | 0.8270± 0.0525 | 0.8270± 0.0525 | 0.8270± 0.0525 | 0.8270± 0.0525 | 0.8270± 0.0525 | 0.8270± 0.0525 |

**Table 9:** Average 5×5 cross validation balanced accuracy and standard deviation on out-of-domain CPTAC test set for the unmodified Gamma-Poisson and Poisson augmentation methods. The scores are reported for different classifier models and different class sizes. The data was augmented given 10% of the original dataset.

| Class Size | Sampling Methods | EBM | KNN | Logistic | RF | SVM-RBF | Average |
| --- | --- | --- | --- | --- | --- | --- | --- |
| max | Gamma-Poisson | 0.6463± 0.0590 | 0.6463± 0.0590 | 0.6463± 0.0590 | 0.6463± 0.0590 | 0.6463± 0.0590 | 0.6463± 0.0590 |
|  | Poisson | 0.6447± 0.0518 | 0.6447± 0.0518 | 0.6447± 0.0518 | 0.6447± 0.0518 | 0.6447± 0.0518 | 0.6447± 0.0518 |
| 500 | Gamma-Poisson | 0.6190± 0.0631 | 0.6190± 0.0631 | 0.6190± 0.0631 | 0.6190± 0.0631 | 0.6190± 0.0631 | 0.6190± 0.0631 |
|  | Poisson | 0.6375± 0.0509 | 0.6375± 0.0509 | 0.6375± 0.0509 | 0.6375± 0.0509 | 0.6375± 0.0509 | 0.6375± 0.0509 |
| 5000 | Gamma-Poisson | 0.6276± 0.0627 | 0.6276± 0.0627 | 0.6276± 0.0627 | 0.6276± 0.0627 | 0.6276± 0.0627 | 0.6276± 0.0627 |
|  | Poisson | 0.6470± 0.0588 | 0.6470± 0.0588 | 0.6470± 0.0588 | 0.6470± 0.0588 | 0.6470± 0.0588 | 0.6470± 0.0588 |

**Table 10:** Average balanced accuracy and standard deviation on in-domain TCGA test set from 5×5 cross validation with different classifier models and an overall average of these models for different augmentation methods. The entire training dataset (100%) was used for augmentation.

| Class Size | Sampling Methods | EBM | KNN | Logistic | RF | SVM-RBF | Average |
| --- | --- | --- | --- | --- | --- | --- | --- |
| Max | Mod. Gamma-Poisson | 0.8841 ± 0.0297 | 0.8331 ± 0.0325 | 0.8936 ± 0.0266 | 0.8782 ± 0.0316 | 0.8896 ± 0.0253 | 0.8757 ± 0.0291 |
| | Inter-Crossover | 0.8948 ± 0.0276 | 0.8465 ± 0.0371 | 0.8917 ± 0.0292 | 0.8821 ± 0.0344 | 0.9011 ± 0.0296 | 0.8832 ± 0.0316 |
| | Intra-Crossover | 0.8796 ± 0.0244 | 0.8434 ± 0.0353 | 0.8837 ± 0.0228 | 0.8763 ± 0.0317 | 0.8808 ± 0.0327 | 0.8728 ± 0.0294 |
| | Mod. Poisson | 0.8819 ± 0.0249 | 0.878 ± 0.0335 | 0.8885 ± 0.0313 | 0.8729 ± 0.0377 | 0.8829 ± 0.0308 | 0.8808 ± 0.0316 |
| | Replacement | 0.8744 ± 0.0328 | 0.8404 ± 0.04 | 0.891 ± 0.022 | 0.8744 ± 0.0288 | 0.8974 ± 0.0253 | 0.8755 ± 0.0298 |
| | SMOTE | 0.8904 ± 0.0225 | 0.8666 ± 0.0342 | 0.8852 ± 0.0306 | 0.8725 ± 0.0306 | 0.8965 ± 0.0243 | 0.8822 ± 0.0284 |
| | Unaugmented | 0.8674 ± 0.0264 | 0.841 ± 0.0354 | 0.8889 ± 0.0319 | 0.8615 ± 0.0335 | 0.8861 ± 0.0315 | 0.869 ± 0.0317 |
| 500 | Mod. Gamma-Poisson | 0.9003 ± 0.0285 | 0.8364 ± 0.0368 | 0.8991 ± 0.0217 | 0.8777 ± 0.0284 | 0.8955 ± 0.0264 | 0.8818 ± 0.0283 |
| | Inter-Crossover | 0.8762 ± 0.0334 | 0.8319 ± 0.041 | 0.8765 ± 0.0266 | 0.8727 ± 0.0275 | 0.8846 ± 0.0314 | 0.8684 ± 0.032 |
| | Intra-Crossover | 0.8827 ± 0.0299 | 0.8485 ± 0.0408 | 0.8763 ± 0.0245 | 0.8704 ± 0.0379 | 0.8873 ± 0.0298 | 0.873 ± 0.0326 |
| | Mod. Poisson | 0.8825 ± 0.027 | 0.8739 ± 0.0402 | 0.8858 ± 0.027 | 0.8651 ± 0.0322 | 0.885 ± 0.0285 | 0.8784 ± 0.031 |
| | Replacement | 0.8684 ± 0.0347 | 0.8226 ± 0.0477 | 0.8792 ± 0.0297 | 0.8728 ± 0.0322 | 0.8887 ± 0.0309 | 0.8663 ± 0.035 |
| | SMOTE | 0.8877 ± 0.0264 | 0.8602 ± 0.0406 | 0.888 ± 0.0315 | 0.8711 ± 0.0324 | 0.8938 ± 0.0261 | 0.8801 ± 0.0314 |
| | Unaugmented | 0.8674 ± 0.0264 | 0.841 ± 0.0354 | 0.8889 ± 0.0319 | 0.8615 ± 0.0335 | 0.8861 ± 0.0315 | 0.869 ± 0.0317 |
| 5000 | Mod. Gamma-Poisson | 0.8927 ± 0.0291 | 0.8583 ± 0.0356 | 0.8955 ± 0.0256 | 0.8781 ± 0.0353 | 0.897 ± 0.0238 | 0.8843 ± 0.0299 |
| | Inter-Crossover | 0.8445 ± 0.0351 | 0.837 ± 0.0403 | 0.8664 ± 0.0298 | 0.8479 ± 0.0378 | 0.8649 ± 0.0297 | 0.8521 ± 0.0345 |
| | Intra-Crossover | 0.8603 ± 0.0281 | 0.8472 ± 0.0465 | 0.8668 ± 0.0278 | 0.8636 ± 0.0377 | 0.8723 ± 0.0257 | 0.8621 ± 0.0332 |
| | Mod. Poisson | 0.8832 ± 0.0282 | 0.8688 ± 0.0383 | 0.8939 ± 0.0276 | 0.8674 ± 0.0352 | 0.8893 ± 0.0272 | 0.8805 ± 0.0313 |
| | Replacement | 0.8654 ± 0.0346 | 0.8048 ± 0.045 | 0.8669 ± 0.0349 | 0.8622 ± 0.0374 | 0.8849 ± 0.0267 | 0.8568 ± 0.0357 |
| | SMOTE | 0.8812 ± 0.033 | 0.8464 ± 0.0405 | 0.8776 ± 0.0296 | 0.8668 ± 0.0346 | 0.8927 ± 0.0292 | 0.8729 ± 0.0334 |
| | Unaugmented | 0.8674 ± 0.0264 | 0.841 ± 0.0354 | 0.8889 ± 0.0319 | 0.8615 ± 0.0335 | 0.8861 ± 0.0315 | 0.869 ± 0.0317 |

**Table 11:** Average balanced accuracy and standard deviation on out-of-domain CPTAC test set from 5×5 cross validation with different classifier models and an overall average of these models for different augmentation methods. The entire training dataset (100%) was used for augmentation.

| Class Size | Sampling Methods | EBM | KNN | Logistic | RF | SVM-RBF | Average |
| --- | --- | --- | --- | --- | --- | --- | --- |
| Max | Mod. Gamma-Poisson | 0.6506 ± 0.0334 | 0.5781 ± 0.0297 | 0.5396 ± 0.0513 | 0.6348 ± 0.0248 | 0.6025 ± 0.029 | 0.6011 ± 0.0336 |
| | Inter-Crossover | 0.67 ± 0.0342 | 0.5876 ± 0.036 | 0.5179 ± 0.0523 | 0.6306 ± 0.0181 | 0.599 ± 0.0268 | 0.601 ± 0.0335 |
| | Intra-Crossover | 0.6474 ± 0.0299 | 0.5947 ± 0.0374 | 0.509 ± 0.0501 | 0.6436 ± 0.0193 | 0.5986 ± 0.0253 | 0.5986 ± 0.0324 |
| | Mod. Poisson | 0.6332 ± 0.0318 | 0.6225 ± 0.026 | 0.537 ± 0.0432 | 0.6309 ± 0.0401 | 0.6013 ± 0.0372 | 0.605 ± 0.0357 |
| | Replacement | 0.6314 ± 0.0331 | 0.5822 ± 0.0311 | 0.55 ± 0.0302 | 0.6369 ± 0.032 | 0.5838 ± 0.0367 | 0.5969 ± 0.0326 |
| | SMOTE | 0.6375 ± 0.0353 | 0.5961 ± 0.0333 | 0.5361 ± 0.0323 | 0.6557 ± 0.0274 | 0.5788 ± 0.0347 | 0.6009 ± 0.0326 |
| | Unaugmented | 0.637 ± 0.0252 | 0.5917 ± 0.0257 | 0.5347 ± 0.0337 | 0.6256 ± 0.0354 | 0.6058 ± 0.0285 | 0.599 ± 0.0297 |
| 500 | Mod. Gamma-Poisson | 0.6732 ± 0.0334 | 0.6 ± 0.0346 | 0.5681 ± 0.0485 | 0.654 ± 0.0196 | 0.6055 ± 0.0313 | 0.6202 ± 0.0335 |
| | Inter-Crossover | 0.7135 ± 0.0242 | 0.6359 ± 0.034 | 0.6213 ± 0.0346 | 0.6657 ± 0.0329 | 0.6303 ± 0.0345 | 0.6534 ± 0.032 |
| | Intra-Crossover | 0.6705 ± 0.0281 | 0.6145 ± 0.0355 | 0.5646 ± 0.0475 | 0.6771 ± 0.0206 | 0.6017 ± 0.0253 | 0.6257 ± 0.0314 |
| | Mod. Poisson | 0.6256 ± 0.0314 | 0.6402 ± 0.03 | 0.5313 ± 0.026 | 0.6353 ± 0.033 | 0.5892 ± 0.0418 | 0.6043 ± 0.0324 |
| | Replacement | 0.6255 ± 0.0261 | 0.5732 ± 0.036 | 0.5531 ± 0.0361 | 0.6275 ± 0.024 | 0.5673 ± 0.0343 | 0.5893 ± 0.0313 |
| | SMOTE | 0.616 ± 0.0343 | 0.6009 ± 0.0378 | 0.5339 ± 0.0268 | 0.6491 ± 0.0309 | 0.5642 ± 0.0344 | 0.5928 ± 0.0328 |
| | Unaugmented | 0.637 ± 0.0252 | 0.5917 ± 0.0257 | 0.5347 ± 0.0337 | 0.6256 ± 0.0354 | 0.6058 ± 0.0285 | 0.599 ± 0.0297 |
| 5000 | Mod. Gamma-Poisson | 0.6845 ± 0.0145 | 0.6253 ± 0.0287 | 0.6344 ± 0.0197 | 0.6775 ± 0.0131 | 0.6243 ± 0.0284 | 0.6492 ± 0.0209 |
| | Inter-Crossover | 0.6532 ± 0.0183 | 0.6748 ± 0.0253 | 0.6533 ± 0.0153 | 0.6821 ± 0.0208 | 0.6592 ± 0.0154 | 0.6645 ± 0.019 |
| | Intra-Crossover | 0.6637 ± 0.0421 | 0.6367 ± 0.0299 | 0.5917 ± 0.038 | 0.673 ± 0.023 | 0.605 ± 0.0279 | 0.634 ± 0.0322 |
| | Mod. Poisson | 0.6243 ± 0.0222 | 0.6354 ± 0.0192 | 0.5293 ± 0.023 | 0.6057 ± 0.0252 | 0.571 ± 0.0358 | 0.5932 ± 0.0251 |
| | Replacement | 0.6271 ± 0.0353 | 0.5725 ± 0.0276 | 0.5588 ± 0.0518 | 0.6206 ± 0.0269 | 0.58 ± 0.0374 | 0.5918 ± 0.0358 |
| | SMOTE | 0.6228 ± 0.0325 | 0.5842 ± 0.0274 | 0.5336 ± 0.0412 | 0.6139 ± 0.0324 | 0.5709 ± 0.041 | 0.5851 ± 0.0349 |
| | Unaugmented | 0.637 ± 0.0252 | 0.5917 ± 0.0257 | 0.5347 ± 0.0337 | 0.6256 ± 0.0354 | 0.6058 ± 0.0285 | 0.599 ± 0.0297 |

**Table 12:** Average balanced accuracy scores and standard deviation on in-domain GSE20713 test set from 5×5 cross-validation for supervised classification of PAM50 subtype prediction with different classifier models and an overall average over these models for the different augmentation methods. Results indicate that augmentation to class size 50 results in Mod. Gamma-Poisson and Mod. Poisson performing significantly better than unaugmented data at significance threshold of 0.05. However, further increasing the class size to 500 results in only the Mod. Poisson performing significantly better than unaugmented data. Best scores in each category are highlighted in bold. Note that the test set size here is ≈ 14 − 15 samples.

| Class Size | Sampling Method | EBM | KNN | Logistic | RF | SVM-RBF | Average |
| --- | --- | --- | --- | --- | --- | --- | --- |
| Max | Mod. Gamma-Poisson | 0.71 ± 0.083 | 0.69 ± 0.101 | 0.66 ± 0.152 | 0.75 ± 0.116 | **0.77 ± 0.095** | 0.72 ± 0.109 |
| | Inter-Crossover | 0.711 ± 0.099 | 0.703 ± 0.096 | **0.692 ± 0.132** | **0.773 ± 0.089** | 0.749 ± 0.097 | 0.726 ± 0.102 |
| | Intra-Crossover | **0.743 ± 0.118** | **0.713 ± 0.089** | 0.671 ± 0.139 | 0.771 ± 0.097 | 0.737 ± 0.100 | 0.727 ± 0.109 |
| | Mod. Poisson | 0.729 ± 0.091 | 0.695 ± 0.107 | 0.671 ± 0.136 | 0.755 ± 0.128 | 0.767 ± 0.086 | 0.723 ± 0.110 |
| | Resampling | 0.735 ± 0.101 | 0.686 ± 0.115 | 0.669 ± 0.130 | 0.769 ± 0.117 | 0.746 ± 0.099 | 0.721 ± 0.113 |
| | SMOTE | 0.739 ± 0.110 | 0.697 ± 0.100 | 0.669 ± 0.131 | 0.743 ± 0.114 | 0.754 ± 0.096 | 0.721 ± 0.110 |
| | Gamma-Poisson | 0.734 ± 0.100 | 0.702 ± 0.119 | 0.686 ± 0.134 | 0.760 ± 0.090 | 0.764 ± 0.100 | 0.729 ± 0.109 |
| | Poisson | 0.731 ± 0.107 | 0.710 ± 0.103 | 0.669 ± 0.128 | 0.757 ± 0.116 | 0.755 ± 0.090 | 0.724 ± 0.109 |
| | Unaugmented | 0.727 ± 0.116 | 0.703 ± 0.100 | 0.663 ± 0.138 | 0.759 ± 0.113 | 0.741 ± 0.107 | 0.719 ± 0.115 |
| 50 | Mod. Gamma-Poisson | **0.731 ± 0.108** | **0.740 ± 0.111** | **0.757 ± 0.112** | **0.763 ± 0.088** | 0.739 ± 0.106 | 0.746 ± 0.105 |
| | Inter-Crossover | 0.656 ± 0.106 | 0.693 ± 0.098 | 0.621 ± 0.113 | 0.665 ± 0.144 | 0.696 ± 0.105 | 0.666 ± 0.113 |
| | Intra-Crossover | 0.712 ± 0.119 | 0.729 ± 0.114 | 0.685 ± 0.133 | 0.757 ± 0.115 | 0.739 ± 0.120 | 0.724 ± 0.120 |
| | Mod. Poisson | 0.728 ± 0.096 | 0.740 ± 0.132 | 0.751 ± 0.129 | 0.751 ± 0.090 | 0.743 ± 0.103 | 0.742 ± 0.110 |
| | Resampling | 0.691 ± 0.114 | 0.685 ± 0.104 | 0.653 ± 0.120 | 0.745 ± 0.122 | 0.709 ± 0.122 | 0.697 ± 0.116 |
| | SMOTE | 0.725 ± 0.072 | 0.689 ± 0.100 | 0.650 ± 0.142 | 0.755 ± 0.111 | 0.706 ± 0.127 | 0.705 ± 0.110 |
| | Gamma-Poisson | 0.712 ± 0.102 | 0.705 ± 0.099 | 0.711 ± 0.116 | 0.715 ± 0.107 | 0.758 ± 0.113 | 0.720 ± 0.107 |
| | Poisson | 0.743 ± 0.117 | 0.728 ± 0.140 | 0.753 ± 0.106 | 0.742 ± 0.116 | **0.772 ± 0.115** | 0.748 ± 0.119 |
| | Unaugmented | 0.727 ± 0.116 | 0.703 ± 0.100 | 0.663 ± 0.138 | 0.759 ± 0.113 | 0.741 ± 0.107 | 0.719 ± 0.115 |
| 500 | Mod. Gamma-Poisson | 0.702 ± 0.109 | 0.737 ± 0.112 | 0.765 ± 0.114 | 0.751 ± 0.094 | 0.737 ± 0.121 | 0.739 ± 0.110 |
| | Inter-Crossover | 0.598 ± 0.094 | 0.687 ± 0.097 | 0.622 ± 0.106 | 0.628 ± 0.104 | 0.682 ± 0.086 | 0.643 ± 0.098 |
| | Intra-Crossover | 0.676 ± 0.108 | 0.745 ± 0.097 | 0.721 ± 0.134 | 0.747 ± 0.118 | 0.740 ± 0.115 | 0.726 ± 0.115 |
| | Mod. Poisson | **0.766 ± 0.091** | 0.750 ± 0.141 | 0.772 ± 0.112 | 0.757 ± 0.101 | **0.751 ± 0.119** | 0.759 ± 0.113 |
| | Resampling | 0.690 ± 0.105 | 0.691 ± 0.112 | 0.639 ± 0.128 | 0.722 ± 0.111 | 0.651 ± 0.117 | 0.679 ± 0.115 |
| | SMOTE | 0.645 ± 0.132 | 0.675 ± 0.103 | 0.634 ± 0.129 | 0.716 ± 0.120 | 0.631 ± 0.127 | 0.660 ± 0.122 |
| | Gamma-Poisson | 0.715 ± 0.107 | **0.755 ± 0.110** | 0.682 ± 0.136 | 0.711 ± 0.111 | 0.720 ± 0.131 | 0.717 ± 0.119 |
| | Poisson | 0.722 ± 0.118 | 0.737 ± 0.120 | **0.783 ± 0.116** | **0.769 ± 0.111** | 0.741 ± 0.114 | 0.750 ± 0.116 |
| | Unaugmented | 0.727 ± 0.116 | 0.703 ± 0.100 | 0.663 ± 0.138 | 0.759 ± 0.113 | 0.741 ± 0.107 | 0.719 ± 0.115 |

**Table 13:** Average balanced accuracy scores and standard deviation on out-of-domain METABRIC test set from 5×5 cross-validation for supervised classification of PAM50 subtype prediction with different classifier models and an overall average over these models for the different augmentation methods. Results indicate that increasing the augmentation size results in only the Mod. Gamma-Poisson and Mod. Poisson methods performing significantly better than unaugmented data at significance threshold of 0.05. Best scores in each category are highlighted in bold.

| Class Size | Sampling Method | EBM | KNN | Logistic | RF | SVM-RBF | Average |
| --- | --- | --- | --- | --- | --- | --- | --- |
| max | Mod. Gamma-Poisson | 0.7175 ± 0.0496 | **0.6677 ± 0.0717** | 0.5809 ± 0.1060 | 0.7146 ± 0.0415 | 0.3370 ± 0.0157 | 0.604 ± 0.0569 |
| | Inter-Crossover | 0.7133 ± 0.0443 | 0.5819 ± 0.0723 | 0.5560 ± 0.1138 | 0.7204 ± 0.0391 | 0.3333 ± 0.0000 | 0.581 ± 0.0539 |
| | Intra-Crossover | 0.7223 ± 0.0286 | 0.6109 ± 0.0624 | 0.5756 ± 0.1138 | 0.7260 ± 0.0309 | 0.3333 ± 0.0000 | 0.594 ± 0.0471 |
| | Mod. Poisson | 0.7138 ± 0.0518 | 0.6470 ± 0.0762 | **0.6192 ± 0.0902** | 0.7090 ± 0.0333 | **0.3423 ± 0.0438** | 0.606 ± 0.0591 |
| | Resampling | 0.7237 ± 0.0384 | 0.5952 ± 0.0670 | 0.5881 ± 0.0970 | 0.7270 ± 0.0327 | 0.3333 ± 0.0000 | 0.593 ± 0.0470 |
| | SMOTE | 0.7388 ± 0.0241 | 0.5856 ± 0.0739 | 0.6011 ± 0.0970 | 0.7121 ± 0.0328 | 0.3333 ± 0.0000 | 0.594 ± 0.0456 |
| | Gamma-Poisson | 0.717 ± 0.054 | 0.685 ± 0.069 | 0.597 ± 0.087 | 0.724 ± 0.032 | 0.342 ± 0.029 | 0.613 ± 0.054 |
| | Poisson | 0.721 ± 0.036 | 0.647 ± 0.084 | 0.584 ± 0.108 | 0.717 ± 0.036 | 0.335 ± 0.006 | 0.601 ± 0.054 |
| | Unaugmented | **0.7412 ± 0.0273** | 0.6195 ± 0.0521 | 0.5923 ± 0.1219 | **0.7361 ± 0.0340** | 0.3333 ± 0.0000 | 0.604 ± 0.0471 |
| 50 | Mod. Gamma-Poisson | 0.7053 ± 0.0640 | **0.7563 ± 0.0343** | 0.6725 ± 0.0756 | 0.7155 ± 0.0573 | 0.7437 ± 0.0532 | 0.719 ± 0.0569 |
| | Inter-Crossover | 0.5005 ± 0.0734 | 0.4607 ± 0.0689 | 0.4509 ± 0.1010 | 0.5532 ± 0.0755 | 0.3333 ± 0.0000 | 0.460 ± 0.0638 |
| | Intra-Crossover | 0.7254 ± 0.0328 | 0.5329 ± 0.0806 | 0.6326 ± 0.1050 | 0.7048 ± 0.0556 | 0.3333 ± 0.0000 | 0.586 ± 0.0548 |
| | Mod. Poisson | 0.7237 ± 0.0416 | 0.7375 ± 0.0299 | 0.6976 ± 0.0602 | 0.7364 ± 0.0400 | **0.7544 ± 0.0580** | 0.730 ± 0.0459 |
| | Resampling | 0.7113 ± 0.0242 | 0.5255 ± 0.0891 | 0.5709 ± 0.1077 | 0.7212 ± 0.0336 | 0.3333 ± 0.0000 | 0.572 ± 0.0509 |
| | SMOTE | 0.7172 ± 0.0263 | 0.5544 ± 0.0813 | 0.5714 ± 0.1082 | 0.7037 ± 0.0412 | 0.3333 ± 0.0000 | 0.576 ± 0.0514 |
| | Gamma-Poisson | 0.662 ± 0.073 | 0.754 ± 0.037 | 0.609 ± 0.105 | 0.714 ± 0.049 | 0.723 ± 0.070 | 0.692 ± 0.067 |
| | Poisson | 0.714 ± 0.057 | 0.747 ± 0.030 | **0.684 ± 0.080** | 0.723 ± 0.058 | 0.750 ± 0.075 | 0.724 ± 0.060 |
| | Unaugmented | **0.7412 ± 0.0273** | 0.6195 ± 0.0521 | 0.5923 ± 0.1219 | **0.7361 ± 0.0340** | 0.3333 ± 0.0000 | 0.604 ± 0.0471 |
| 500 | Mod. Gamma-Poisson | 0.7312 ± 0.0440 | **0.7536 ± 0.0331** | 0.7697 ± 0.0429 | 0.7237 ± 0.0519 | 0.7600 ± 0.0540 | 0.748 ± 0.0452 |
| | Inter-Crossover | 0.4097 ± 0.0515 | 0.4100 ± 0.0627 | 0.4264 ± 0.0845 | 0.4273 ± 0.0656 | 0.3333 ± 0.0000 | 0.401 ± 0.0529 |
| | Intra-Crossover | 0.7207 ± 0.0362 | 0.4704 ± 0.0703 | 0.6048 ± 0.0858 | 0.5403 ± 0.0839 | 0.3333 ± 0.0000 | 0.534 ± 0.0552 |
| | Mod. Poisson | 0.7308 ± 0.0359 | 0.7471 ± 0.0345 | 0.7644 ± 0.0519 | **0.7405 ± 0.0386** | 0.7553 ± 0.0521 | 0.748 ± 0.0426 |
| | Resampling | 0.6956 ± 0.0423 | 0.5217 ± 0.0748 | 0.5517 ± 0.0927 | 0.7134 ± 0.0340 | 0.3333 ± 0.0000 | 0.563 ± 0.0488 |
| | SMOTE | 0.6883 ± 0.0534 | 0.5183 ± 0.0776 | 0.5644 ± 0.0880 | 0.6578 ± 0.0557 | 0.3333 ± 0.0000 | 0.552 ± 0.0549 |
| | Gamma-Poisson | 0.681 ± 0.067 | 0.753 ± 0.034 | 0.635 ± 0.101 | 0.664 ± 0.075 | 0.658 ± 0.087 | 0.678 ± 0.073 |
| | Poisson | 0.739 ± 0.051 | 0.749 ± 0.029 | **0.775 ± 0.045** | 0.732 ± 0.057 | **0.770 ± 0.037** | 0.753 ± 0.044 |
| | Unaugmented | **0.7412 ± 0.0273** | 0.6195 ± 0.0521 | 0.5923 ± 0.1219 | 0.7361 ± 0.0340 | 0.3333 ± 0.0000 | 0.604 ± 0.0471 |

**Table 14:** Average balanced accuracy and standard deviation on in-domain TCGA test set from 5×5 cross validation with different classifier models and an overall average of these models for different augmentation methods. A portion of the training dataset (10%) was used for augmentation.

| Class Size | Sampling Methods | EBM | KNN | Logistic | RF | SVM-RBF | Average |
| --- | --- | --- | --- | --- | --- | --- | --- |
| Max | Mod. Gamma-Poisson | $0.8126 \pm 0.0455$ | $0.7525 \pm 0.0542$ | $0.8363 \pm 0.0609$ | $0.7914 \pm 0.0684$ | $0.8182 \pm 0.0567$ | $0.8022 \pm 0.0572$ |
| | Inter-Crossover | $0.8092 \pm 0.0536$ | $0.7666 \pm 0.0463$ | $0.8349 \pm 0.058$ | $0.8129 \pm 0.0433$ | $0.8258 \pm 0.0581$ | $0.8099 \pm 0.0518$ |
| | Intra-Crossover | $0.8141 \pm 0.0524$ | $0.7697 \pm 0.0535$ | $0.8272 \pm 0.0589$ | $0.7973 \pm 0.0568$ | $0.822 \pm 0.0554$ | $0.8061 \pm 0.0554$ |
| | Mod. Poisson | $0.8049 \pm 0.0593$ | $0.8191 \pm 0.0418$ | $0.8323 \pm 0.0564$ | $0.7838 \pm 0.0682$ | $0.7965 \pm 0.0593$ | $0.8073 \pm 0.057$ |
| | Replacement | $0.7983 \pm 0.052$ | $0.7467 \pm 0.0628$ | $0.836 \pm 0.0566$ | $0.7842 \pm 0.0533$ | $0.8097 \pm 0.0563$ | $0.795 \pm 0.0562$ |
| | SMOTE | $0.806 \pm 0.0513$ | $0.7827 \pm 0.0534$ | $0.835 \pm 0.0559$ | $0.7918 \pm 0.0659$ | $0.8203 \pm 0.0562$ | $0.8072 \pm 0.0566$ |
| | Unaugmented | $0.7707 \pm 0.0551$ | $0.7494 \pm 0.062$ | $0.8165 \pm 0.0561$ | $0.7858 \pm 0.047$ | $0.7863 \pm 0.0509$ | $0.7817 \pm 0.0542$ |
| 500 | Mod. Gamma-Poisson | $0.8341 \pm 0.0497$ | $0.8012 \pm 0.0521$ | $0.8433 \pm 0.0498$ | $0.8023 \pm 0.0633$ | $0.8412 \pm 0.0492$ | $0.8244 \pm 0.0528$ |
| | Inter-Crossover | $0.789 \pm 0.051$ | $0.7889 \pm 0.0414$ | $0.7867 \pm 0.0522$ | $0.7851 \pm 0.0483$ | $0.8116 \pm 0.0483$ | $0.7923 \pm 0.0482$ |
| | Intra-Crossover | $0.7824 \pm 0.0479$ | $0.7856 \pm 0.0519$ | $0.7974 \pm 0.0488$ | $0.7521 \pm 0.0571$ | $0.7849 \pm 0.0519$ | $0.7805 \pm 0.0515$ |
| | Mod. Poisson | $0.822 \pm 0.0589$ | $0.8076 \pm 0.0542$ | $0.8463 \pm 0.0449$ | $0.7724 \pm 0.0533$ | $0.6977 \pm 0.0999$ | $0.7892 \pm 0.0622$ |
| | Replacement | $0.8009 \pm 0.0527$ | $0.7399 \pm 0.0625$ | $0.8336 \pm 0.0521$ | $0.7653 \pm 0.0572$ | $0.8118 \pm 0.0614$ | $0.7903 \pm 0.0572$ |
| | SMOTE | $0.8209 \pm 0.0568$ | $0.8036 \pm 0.0577$ | $0.8403 \pm 0.0597$ | $0.7825 \pm 0.0623$ | $0.8232 \pm 0.0553$ | $0.8141 \pm 0.0584$ |
| | Unaugmented | $0.7707 \pm 0.0551$ | $0.7494 \pm 0.062$ | $0.8165 \pm 0.0561$ | $0.7858 \pm 0.047$ | $0.7863 \pm 0.0509$ | $0.7817 \pm 0.0542$ |
| 5000 | Mod. Gamma-Poisson | $0.838 \pm 0.0521$ | $0.8154 \pm 0.0505$ | $0.8353 \pm 0.0618$ | $0.7972 \pm 0.0661$ | $0.841 \pm 0.055$ | $0.8254 \pm 0.0571$ |
| | Inter-Crossover | $0.761 \pm 0.0529$ | $0.7852 \pm 0.0543$ | $0.7574 \pm 0.0516$ | $0.7473 \pm 0.0553$ | $0.7615 \pm 0.0557$ | $0.7625 \pm 0.054$ |
| | Intra-Crossover | $0.7661 \pm 0.0505$ | $0.7474 \pm 0.0514$ | $0.7785 \pm 0.057$ | $0.7006 \pm 0.0663$ | $0.7672 \pm 0.04$ | $0.752 \pm 0.053$ |
| | Mod. Poisson | $0.8221 \pm 0.054$ | $0.7969 \pm 0.0548$ | $0.8381 \pm 0.046$ | $0.7742 \pm 0.0508$ | $0.6887 \pm 0.1054$ | $0.784 \pm 0.0622$ |
| | Replacement | $0.801 \pm 0.0569$ | $0.7407 \pm 0.063$ | $0.8273 \pm 0.0506$ | $0.7688 \pm 0.0631$ | $0.8119 \pm 0.0622$ | $0.7899 \pm 0.0591$ |
| | SMOTE | $0.8157 \pm 0.0562$ | $0.7994 \pm 0.0571$ | $0.835 \pm 0.0556$ | $0.7693 \pm 0.0581$ | $0.823 \pm 0.0567$ | $0.8085 \pm 0.0568$ |
| | Unaugmented | $0.7707 \pm 0.0551$ | $0.7494 \pm 0.062$ | $0.8165 \pm 0.0561$ | $0.7858 \pm 0.047$ | $0.7863 \pm 0.0509$ | $0.7817 \pm 0.0542$ |

**Table 15:**
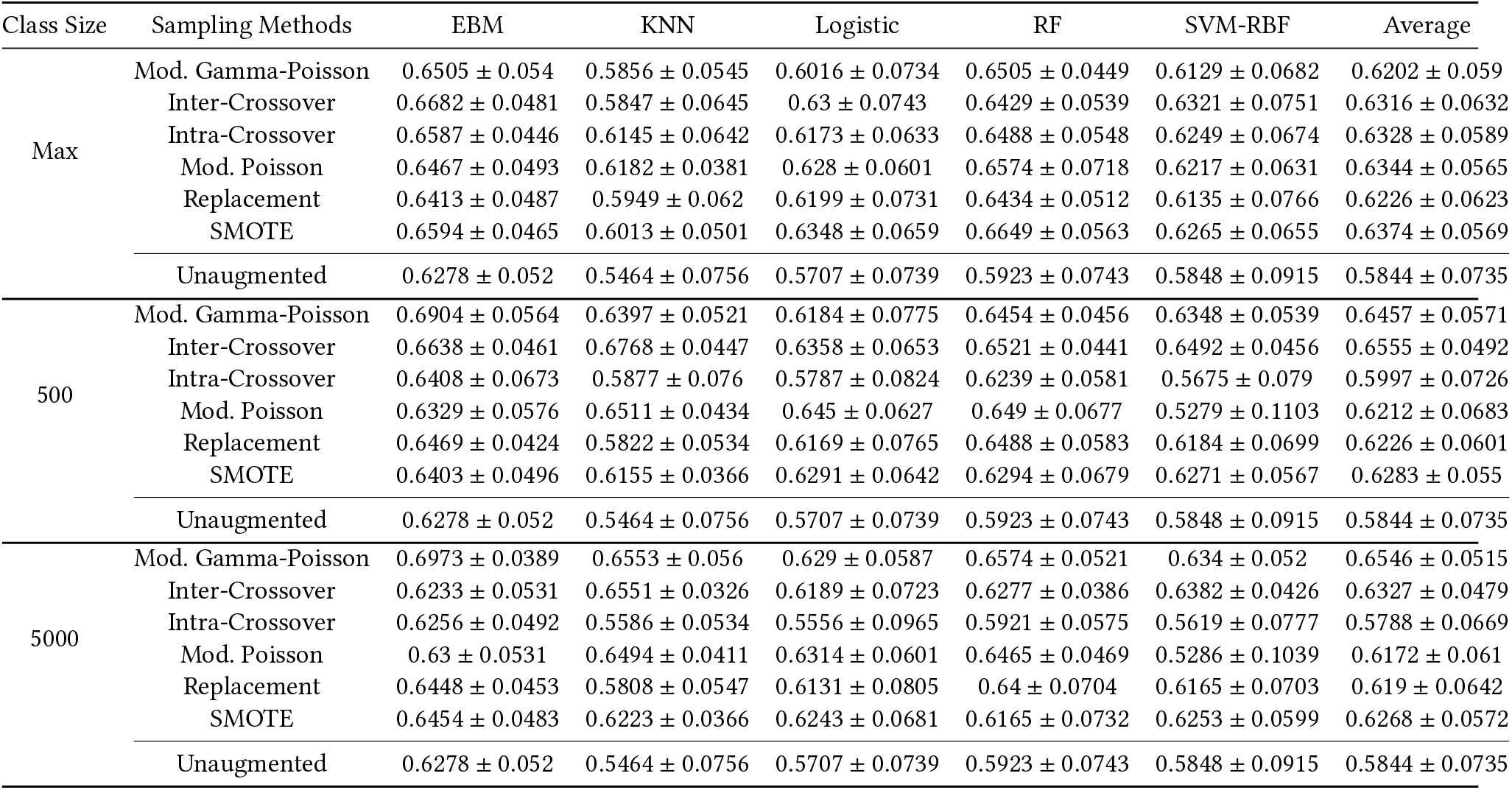
Average balanced accuracy and standard deviation on out-of-domain CPTAC test set from 5×5 cross validation with different classifier models and an overall average of these models for different augmentation methods. A portion of the training dataset (10%) was used for augmentation.

| Class Size | Sampling Methods | EBM | KNN | Logistic | RF | SVM-RBF | Average |
| --- | --- | --- | --- | --- | --- | --- | --- |
| Max | Mod. Gamma-Poisson | 0.6505 ± 0.054 | 0.5856 ± 0.0545 | 0.6016 ± 0.0734 | 0.6505 ± 0.0449 | 0.6129 ± 0.0682 | 0.6202 ± 0.059 |
| | Inter-Crossover | 0.6682 ± 0.0481 | 0.5847 ± 0.0645 | 0.63 ± 0.0743 | 0.6429 ± 0.0539 | 0.6321 ± 0.0751 | 0.6316 ± 0.0632 |
| | Intra-Crossover | 0.6587 ± 0.0446 | 0.6145 ± 0.0642 | 0.6173 ± 0.0633 | 0.6488 ± 0.0548 | 0.6249 ± 0.0674 | 0.6328 ± 0.0589 |
| | Mod. Poisson | 0.6467 ± 0.0493 | 0.6182 ± 0.0381 | 0.628 ± 0.0601 | 0.6574 ± 0.0718 | 0.6217 ± 0.0631 | 0.6344 ± 0.0565 |
| | Replacement | 0.6413 ± 0.0487 | 0.5949 ± 0.062 | 0.6199 ± 0.0731 | 0.6434 ± 0.0512 | 0.6135 ± 0.0766 | 0.6226 ± 0.0623 |
| | SMOTE | 0.6594 ± 0.0465 | 0.6013 ± 0.0501 | 0.6348 ± 0.0659 | 0.6649 ± 0.0563 | 0.6265 ± 0.0655 | 0.6374 ± 0.0569 |
| | Unaugmented | 0.6278 ± 0.052 | 0.5464 ± 0.0756 | 0.5707 ± 0.0739 | 0.5923 ± 0.0743 | 0.5848 ± 0.0915 | 0.5844 ± 0.0735 |
| 500 | Mod. Gamma-Poisson | 0.6904 ± 0.0564 | 0.6397 ± 0.0521 | 0.6184 ± 0.0775 | 0.6454 ± 0.0456 | 0.6348 ± 0.0539 | 0.6457 ± 0.0571 |
| | Inter-Crossover | 0.6638 ± 0.0461 | 0.6768 ± 0.0447 | 0.6358 ± 0.0653 | 0.6521 ± 0.0441 | 0.6492 ± 0.0456 | 0.6555 ± 0.0492 |
| | Intra-Crossover | 0.6408 ± 0.0673 | 0.5877 ± 0.076 | 0.5787 ± 0.0824 | 0.6239 ± 0.0581 | 0.5675 ± 0.079 | 0.5997 ± 0.0726 |
| | Mod. Poisson | 0.6329 ± 0.0576 | 0.6511 ± 0.0434 | 0.645 ± 0.0627 | 0.649 ± 0.0677 | 0.5279 ± 0.1103 | 0.6212 ± 0.0683 |
| | Replacement | 0.6469 ± 0.0424 | 0.5822 ± 0.0534 | 0.6169 ± 0.0765 | 0.6488 ± 0.0583 | 0.6184 ± 0.0699 | 0.6226 ± 0.0601 |
| | SMOTE | 0.6403 ± 0.0496 | 0.6155 ± 0.0366 | 0.6291 ± 0.0642 | 0.6294 ± 0.0679 | 0.6271 ± 0.0567 | 0.6283 ± 0.055 |
| | Unaugmented | 0.6278 ± 0.052 | 0.5464 ± 0.0756 | 0.5707 ± 0.0739 | 0.5923 ± 0.0743 | 0.5848 ± 0.0915 | 0.5844 ± 0.0735 |
| 5000 | Mod. Gamma-Poisson | 0.6973 ± 0.0389 | 0.6553 ± 0.056 | 0.629 ± 0.0587 | 0.6574 ± 0.0521 | 0.634 ± 0.052 | 0.6546 ± 0.0515 |
| | Inter-Crossover | 0.6233 ± 0.0531 | 0.6551 ± 0.0326 | 0.6189 ± 0.0723 | 0.6277 ± 0.0386 | 0.6382 ± 0.0426 | 0.6327 ± 0.0479 |
| | Intra-Crossover | 0.6256 ± 0.0492 | 0.5586 ± 0.0534 | 0.5556 ± 0.0965 | 0.5921 ± 0.0575 | 0.5619 ± 0.0777 | 0.5788 ± 0.0669 |
| | Mod. Poisson | 0.63 ± 0.0531 | 0.6494 ± 0.0411 | 0.6314 ± 0.0601 | 0.6465 ± 0.0469 | 0.5286 ± 0.1039 | 0.6172 ± 0.061 |
| | Replacement | 0.6448 ± 0.0453 | 0.5808 ± 0.0547 | 0.6131 ± 0.0805 | 0.64 ± 0.0704 | 0.6165 ± 0.0703 | 0.619 ± 0.0642 |
| | SMOTE | 0.6454 ± 0.0483 | 0.6223 ± 0.0366 | 0.6243 ± 0.0681 | 0.6165 ± 0.0732 | 0.6253 ± 0.0599 | 0.6268 ± 0.0572 |
| | Unaugmented | 0.6278 ± 0.052 | 0.5464 ± 0.0756 | 0.5707 ± 0.0739 | 0.5923 ± 0.0743 | 0.5848 ± 0.0915 | 0.5844 ± 0.0735 |

**Table 16:** Average balanced accuracy and standard deviation of different classifier models on VAE generated samples modeled on in-domain TCGA test set from 5×5 cross validation and an overall average of these models for different augmentation methods. A portion of the real training dataset (10%) was used for augmentation and each class was augmented to size 500.

| Sampling Methods | EBM | KNN | Logistic | RF | SVM-RBF | Average |
| --- | --- | --- | --- | --- | --- | --- |
| Mod. Gamma-Poisson | 0.7794 ± 0.0548 | 0.7694 ± 0.0504 | 0.7829 ± 0.0565 | 0.7689 ± 0.0553 | 0.7851 ± 0.0508 | 0.7771 ± 0.0536 |
| Inter-Crossover | 0.749 ± 0.0558 | 0.7362 ± 0.058 | 0.7609 ± 0.0515 | 0.7524 ± 0.0538 | 0.7619 ± 0.0465 | 0.7521 ± 0.0531 |
| Intra-Crossover | 0.76 ± 0.0524 | 0.7568 ± 0.0526 | 0.7877 ± 0.055 | 0.7677 ± 0.0561 | 0.7763 ± 0.0578 | 0.7697 ± 0.0548 |
| Mod. Poisson | 0.7322 ± 0.0714 | 0.7314 ± 0.0611 | 0.7327 ± 0.0626 | 0.7237 ± 0.0839 | 0.7255 ± 0.0681 | 0.7291 ± 0.0694 |
| Replacement | 0.7612 ± 0.0516 | 0.7396 ± 0.0687 | 0.7719 ± 0.0565 | 0.7567 ± 0.0561 | 0.7623 ± 0.0573 | 0.7583 ± 0.0580 |
| SMOTE | 0.7639 ± 0.0629 | 0.7468 ± 0.0528 | 0.7619 ± 0.0599 | 0.7544 ± 0.0569 | 0.7585 ± 0.0548 | 0.7571 ± 0.0575 |
| Unaugmented | 0.2517 ± 0.0091 | 0.2502 ± 0.0122 | 0.2514 ± 0.0209 | 0.2504 ± 0.0162 | 0.25 ± 0.0 | 0.2507 ± 0.0117 |

**Table 17:** Average balanced accuracy and standard deviation of different classifier models on VAE generated samples modeled on out-of-domain CPTAC test set from 5×5 cross validation and an overall average of these models for different augmentation methods. A portion of the real training dataset (10%) was used for augmentation and each class was augmented to size 500.

| Sampling Methods | EBM | KNN | Logistic | RF | SVM-RBF | Average |
| --- | --- | --- | --- | --- | --- | --- |
| Mod. Gamma-Poisson | 0.6223 ± 0.0574 | 0.6018 ± 0.0785 | 0.6009 ± 0.059 | 0.5978 ± 0.0653 | 0.6157 ± 0.0704 | 0.6077 ± 0.0661 |
| Inter-Crossover | 0.649 ± 0.0632 | 0.6719 ± 0.0657 | 0.67 ± 0.0622 | 0.6489 ± 0.0602 | 0.678 ± 0.0635 | 0.6636 ± 0.0630 |
| Intra-Crossover | 0.6373 ± 0.0754 | 0.638 ± 0.0585 | 0.6502 ± 0.0624 | 0.6225 ± 0.0675 | 0.6448 ± 0.064 | 0.6386 ± 0.0656 |
| Mod. Poisson | 0.6147 ± 0.066 | 0.6201 ± 0.0657 | 0.6305 ± 0.0628 | 0.5892 ± 0.0621 | 0.6164 ± 0.0569 | 0.6142 ± 0.0627 |
| Replacement | 0.6198 ± 0.0653 | 0.5848 ± 0.0708 | 0.6254 ± 0.0525 | 0.6014 ± 0.0739 | 0.6147 ± 0.0568 | 0.6092 ± 0.0649 |
| SMOTE | 0.6203 ± 0.0546 | 0.617 ± 0.0631 | 0.624 ± 0.0528 | 0.6094 ± 0.0616 | 0.6209 ± 0.0583 | 0.6183 ± 0.0581 |
| Unaugmented | 0.2573 ± 0.0153 | 0.2512 ± 0.0069 | 0.2524 ± 0.0213 | 0.2536 ± 0.0162 | 0.25 ± 0.0 | 0.2529 ± 0.0119 |

**Table 18:** Average balanced accuracy and standard deviation on in-domain TCGA test data of different classifier models in predicting the MSI status from embeddings retrieved from a VAE and an overall average of these models for different augmentation methods. A portion of the real training dataset (10%) was used for augmentation and each class was augmented to size 500.

| Sampling Methods | EBM | KNN | Logistic | RF | SVM-RBF | Average |
| --- | --- | --- | --- | --- | --- | --- |
| Mod. Gamma-Poisson | 0.5723 ± 0.0483 | 0.5958 ± 0.0472 | 0.6117 ± 0.0597 | 0.5895 ± 0.0617 | 0.5865 ± 0.0713 | 0.5912 ± 0.0576 |
| Inter-Crossover | 0.5683 ± 0.0423 | 0.6184 ± 0.0498 | 0.6216 ± 0.0667 | 0.5962 ± 0.0643 | 0.6187 ± 0.0578 | 0.6046 ± 0.0562 |
| Intra-Crossover | 0.5667 ± 0.0538 | 0.6177 ± 0.0534 | 0.6222 ± 0.0691 | 0.5983 ± 0.0587 | 0.6223 ± 0.062 | 0.6054 ± 0.0594 |
| Mod. Poisson | 0.5736 ± 0.0601 | 0.5879 ± 0.0677 | 0.6023 ± 0.0615 | 0.5887 ± 0.0472 | 0.5772 ± 0.0555 | 0.5859 ± 0.0584 |
| Replacement | 0.6057 ± 0.0572 | 0.5872 ± 0.0543 | 0.6391 ± 0.0482 | 0.619 ± 0.0467 | 0.6124 ± 0.06 | 0.6127 ± 0.0533 |
| SMOTE | 0.5875 ± 0.0542 | 0.5707 ± 0.0548 | 0.6229 ± 0.0598 | 0.6149 ± 0.0591 | 0.5945 ± 0.062 | 0.5981 ± 0.0580 |
| Unaugmented | 0.4873 ± 0.0385 | 0.483 ± 0.0349 | 0.484 ± 0.04 | 0.4862 ± 0.0406 | 0.4932 ± 0.0191 | 0.4867 ± 0.0346 |

**Table 19:** Average balanced accuracy and standard deviation on out-of-domain CPTAC test data of different classifier models in predicting the MSI status from embeddings retrieved from a VAE and an overall average of these models for different augmentation methods. A portion of the real training dataset (10%) was used for augmentation and each class was augmented to size 500.

| Sampling Methods | EBM | KNN | Logistic | RF | SVM-RBF | Average |
| --- | --- | --- | --- | --- | --- | --- |
| Mod. Gamma-Poisson | 0.5808 ± 0.0826 | 0.6444 ± 0.0957 | 0.6823 ± 0.1153 | 0.6208 ± 0.0896 | 0.6458 ± 0.1118 | 0.6348 ± 0.0990 |
| Inter-Crossover | 0.6601 ± 0.1057 | 0.7332 ± 0.1307 | 0.7087 ± 0.1329 | 0.6897 ± 0.104 | 0.728 ± 0.1256 | 0.7039 ± 0.1198 |
| Intra-Crossover | 0.6394 ± 0.0859 | 0.7097 ± 0.1083 | 0.7333 ± 0.122 | 0.7016 ± 0.1163 | 0.7592 ± 0.1111 | 0.7086 ± 0.1087 |
| Mod. Poisson | 0.6345 ± 0.0792 | 0.6534 ± 0.1046 | 0.6996 ± 0.107 | 0.668 ± 0.0658 | 0.6463 ± 0.097 | 0.6604 ± 0.0907 |
| Replacement | 0.6495 ± 0.0701 | 0.637 ± 0.1028 | 0.7616 ± 0.1182 | 0.7021 ± 0.1014 | 0.6915 ± 0.1133 | 0.6883 ± 0.1012 |
| SMOTE | 0.6152 ± 0.0916 | 0.5956 ± 0.1044 | 0.6992 ± 0.1062 | 0.6892 ± 0.0947 | 0.6456 ± 0.089 | 0.6489 ± 0.0972 |
| Unaugmented | 0.5002 ± 0.0274 | 0.5144 ± 0.0411 | 0.4903 ± 0.0425 | 0.5078 ± 0.0374 | 0.4989 ± 0.0265 | 0.5023 ± 0.0350 |

**Table 20:** Average balanced accuracy and standard deviation on in-domain TCGA test data of different classifier models in predicting the CIMP status from embeddings retrieved from a VAE and an overall average of these models for different augmentation methods. A portion of the real training dataset (10%) was used for augmentation and each class was augmented to size 500.

| Sampling Methods | EBM | KNN | Logistic | RF | SVM-RBF | Average |
| --- | --- | --- | --- | --- | --- | --- |
| Mod. Gamma-Poisson | 0.4329 ± 0.0757 | 0.5189 ± 0.0562 | 0.5275 ± 0.0344 | 0.5153 ± 0.0572 | 0.4867 ± 0.062 | 0.4963 ± 0.0571 |
| Inter-Crossover | 0.4606 ± 0.0789 | 0.5633 ± 0.0572 | 0.5565 ± 0.0489 | 0.5183 ± 0.0597 | 0.5156 ± 0.0671 | 0.5229 ± 0.0624 |
| Intra-Crossover | 0.4538 ± 0.078 | 0.5358 ± 0.0571 | 0.5451 ± 0.0476 | 0.5085 ± 0.0688 | 0.5076 ± 0.0624 | 0.5102 ± 0.0628 |
| Mod. Poisson | 0.4418 ± 0.0718 | 0.5024 ± 0.0705 | 0.5302 ± 0.0538 | 0.531 ± 0.0763 | 0.4818 ± 0.0797 | 0.4974 ± 0.0704 |
| Replacement | 0.4502 ± 0.0639 | 0.514 ± 0.0661 | 0.542 ± 0.0526 | 0.5288 ± 0.0801 | 0.5127 ± 0.0645 | 0.5095 ± 0.0654 |
| SMOTE | 0.4595 ± 0.0902 | 0.4968 ± 0.0615 | 0.5441 ± 0.0559 | 0.5308 ± 0.0699 | 0.5018 ± 0.0779 | 0.5066 ± 0.0711 |
| Unaugmented | 0.3452 ± 0.0392 | 0.3429 ± 0.0595 | 0.3363 ± 0.032 | 0.3411 ± 0.0646 | 0.3386 ± 0.0183 | 0.3408 ± 0.0427 |

### H.2 Classification performance

*Non-overlapping signatures*. Table 10 describes the results of CMS classification of the augmentation methods on in-domain TCGA test data. Figure 12 and Figure 13 illustrate the distribution of the ROC-AUC scores for the same task across CV splits for in-domain and out-of-domain test data, respectively. From Table 10, we observe that the Mod. Gamma-Poisson augmentation method is best suited for in-domain data. At class size Max, the average balanced accuracy is 87.57% ± 0.029, at class size 500 it increases to 88.18% ± 0.028, and it begins to saturate at class size 5000 with 88.43% ± 0.029. As expected, the choice of class size impacts the effectiveness of the data augmentation methods. Interestingly, the distribution-based Mod. Gamma-Poisson perfoms best when the number of augmented samples is pushed to the extreme, benefiting from the increased class size in contrast to other methods. However, the trend indicates that over-augmentation offers little to no benefits at the expense of increased computation (≈ 0.3% performance gain from class size 500 to 5000).

#### H.2.2 Overlapping signatures

Figure 14 illustrates the distribution of balanced accuracy scores for PAM50 subtype classification across GSE20713 CV test splits. Figure 15 and Figure 16 illustrate the distribution of the ROC-AUC scores across CV splits for in-domain and out-of-domain test data, respectively. Table 12 and Table 13 describe the average balanced accuracy scores on in-domain GSE20713 and out-of-domain METABRIC test sets for each classifier and augmentation method. At lower augmentation sizes, both the modified and unmodified Poisson sampling methods are also associated with best performance. The negligible performance differences between these two methods could be attributed to the low number of both real samples and the number of generated samples, which limits the diversity of the generated samples by the Mod. Poisson sampling method. Although the Mod. Poisson sampling method sometimes performs on-par, particularly in the lower augmentation size regime, the Mod. Gamma-Poisson should be the preferred method as it can handle over-dispersion.

### H.3 Effect of 10% real data

Results on classification performance when only 10% of the real data are provided is shown in Table 14 for in-domain testing and Table 15 for out-of-domain testing.

### H.4 VAE generated sample classification

Figure 17 illustrates the distribution of balanced accuracy of CMS classification on VAE generated samples modeled on TCGA test data and Table 16 quantifies the average balanced accuracy for each classification model and augmentation method. Similarly, Table 17 quantifies the average balanced accuracy for each classification model and augmentation method for VAE generated samples modeled on CPTAC test data. As expected from in-domain testing, augmentation is absolutely necessary in this low-data regime to learn anything useful from the data. On average (see Figure 17), almost all methods are on-par with each other, with an average difference of around 2% between the methods. The Mod. Gamma-Poisson augmentation method achieves the best performances with the classification models, with an average accuracy of 77.71% ± 0.0536. All methods are significantly better than the unaugmented case.

### H.5 Predicting clinical variables

Table 18 and Table 19 show results for in-domain TCGA test data and out-of-domain CPTAC data, respectively, of different classifier models in predicting the MSI status from VAE embeddings quantified as average balanced accuracy. Table 20 shows the average balanced accuracy on in-domain TCGA test data of different classifier models in predicting the CIMP status from VAE embeddings.

## Notes

https://github.com/PaccMann/transcriptomic_signature_sampling

### Competing Interest Statement

The authors have declared no competing interest.

### Summary of Updates

The paper has undergone restructuring to better highlight our contributions to both signature-dependent and signature-independent sampling. This includes new experiments on breast cancer to consider the scenario wherein signatures have overlapping genes. Additionally, we have also included experiments to analyse the behaviour of augmentation methods in cases of increasing percentages of available data (i.e available data is limited).

## References

[1] James H Bullard, Elizabeth Purdom, Kasper D Hansen, and Sandrine Dudoit. Evaluation of statistical methods for normalization and differential expression in mrna-seq experiments. BMC bioinformatics, 11(1):1–13, 2010.

[2] Likun Wang, Zhixing Feng, Xi Wang, Xiaowo Wang, and Xuegong Zhang. Degseq: an r package for identifying differentially expressed genes from rna-seq data. Bioinformatics, 26(1):136–138, 2010.

[3] Michael I Love, Wolfgang Huber, and Simon Anders. Moderated estimation of fold change and dispersion for rna-seq data with deseq2. Genome biology, 15(12):1–21, 2014.

[4] Mark D Robinson, Davis J McCarthy, and Gordon K Smyth. edger: a bioconductor package for differential expression analysis of digital gene expression data. bioinformatics, 26(1):139–140, 2010.

[5] Dan Hendrycks and Kevin Gimpel. A baseline for detecting misclassified and out-of-distribution examples in neural networks. arXiv preprint 1610.02136, 2016.

[6] Ian Goodfellow, Jean Pouget-Abadie, Mehdi Mirza, Bing Xu, David Warde-Farley, Sherjil Ozair, Aaron Courville, and Yoshua Bengio. Generative adversarial networks. Communications of the ACM, 63(11):139–144, 2020.

[7] Ramon Viñas, Helena Andrés-Terré, Pietro Liò, and Kevin Bryson. Adversarial generation of gene expression data. Bioinformatics, 38(3):730–737, 2022.

[8] Poonam Chaudhari, Himanshu Agrawal, and Ketan Kotecha. Data augmentation using mg-gan for improved cancer classification on gene expression data. Soft Computing, 24(15):11381–11391, 2020.

[9] Ozlem Yersal and Sabri Barutca. Biological subtypes of breast cancer: Prognostic and therapeutic implications. World journal of clinical oncology, 5(3):412, 2014.

[10] Nuria Rodriguez-Salas, Gema Dominguez, Rodrigo Barderas, Marta Mendiola, Xabier García-Albéniz, Juan Maurel, and Jaime Feliu Batlle. Clinical relevance of colorectal cancer molecular subtypes. Critical reviews in oncology/hematology, 109:9–19, 2017.

[11] Xu Zhou, Kai Hu, Peter Bailey, Christoph Springfeld, Susanne Roth, Roma Kurilov, Benedikt Brors, Thomas Gress, Malte Buchholz, Jingyu An, et al. Clinical impact of molecular subtyping of pancreatic cancer. Frontiers in cell and developmental biology, 9:743908, 2021.

[12] Justin Guinney, Rodrigo Dienstmann, Xin Wang, Aurélien De Reynies, Andreas Schlicker, Charlotte Soneson, Laetitia Marisa, Paul Roepman, Gift Nyamundanda, Paolo Angelino, et al. The consensus molecular subtypes of colorectal cancer. Nature medicine, 21(11):1350–1356, 2015.

[13] Steven A Buechler, Melissa T Stephens, Amanda B Hummon, Katelyn Ludwig, Emily Cannon, Tonia C Carter, Jeffrey Resnick, Yesim Gökmen-Polar, and Sunil S Badve. Colotype: a forty gene signature for consensus molecular subtyping of colorectal cancer tumors using whole-genome assay or targeted rna-sequencing. Scientific reports, 10(1):1–13, 2020.

[14] Major Greenwood and G Udny Yule. An inquiry into the nature of frequency distributions representative of multiple happenings with particular reference to the occurrence of multiple attacks of disease or of repeated accidents. Journal of the Royal statistical society, 83(2):255–279, 1920.

[15] Tim Barry. Tim barry: Gamma, poisson, and negative binomial distributions, 2020.

[16] Cancer Genome Atlas Network et al. Comprehensive molecular characterization of human colon and rectal cancer. Nature, 487(7407):330, 2012.

[17] Nathan J Edwards, Mauricio Oberti, Ratna R Thangudu, Shuang Cai, Peter B McGarvey, Shine Jacob, Subha Madhavan, and Karen A Ketchum. The cptac data portal: a resource for cancer proteomics research. Journal of proteome research, 14(6):2707–2713, 2015.

[18] Sarah Dedeurwaerder, Christine Desmedt, Emilie Calonne, Sandeep K Singhal, Benjamin Haibe-Kains, Matthieu Defrance, Stefan Michiels, Michael Volkmar, Rachel Deplus, Judith Luciani, et al. Dna methylation profiling reveals a predominant immune component in breast cancers. EMBO molecular medicine, 3(12):726–741, 2011.

[19] Christina Curtis, Sohrab P Shah, Suet-Feung Chin, Gulisa Turashvili, Oscar M Rueda, Mark J Dunning, Doug Speed, Andy G Lynch, Shamith Samarajiwa, Yinyin Yuan, et al. The genomic and transcriptomic architecture of 2,000 breast tumours reveals novel subgroups. Nature, 486(7403):346–352, 2012.

[20] Bernard Pereira, Suet-Feung Chin, Oscar M Rueda, Hans-Kristian Moen Vollan, Elena Provenzano, Helen A Bardwell, Michelle Pugh, Linda Jones, Roslin Russell, Stephen-John Sammut, et al. The somatic mutation profiles of 2,433 breast cancers refine their genomic and transcriptomic landscapes. Nature communications, 7(1):1–16, 2016.

[21] Guillaume Lemaître, Fernando Nogueira, and Christos K. Aridas. Imbalancedlearn: A python toolbox to tackle the curse of imbalanced datasets in machine learning. Journal of Machine Learning Research, 18(17):1–5, 2017.

[22] Ana Conesa, Pedro Madrigal, Sonia Tarazona, David Gomez-Cabrero, Alejandra Cervera, Andrew McPherson, Michal Wojciech Szczesniak, Daniel J Gaffney, Laura L Elo, Xuegong Zhang, et al. A survey of best practices for rna-seq data analysis. Genome biology, 17(1):1–19, 2016.

[23] Diederik P Kingma and Max Welling. Auto-encoding variational bayes. In Proceedings of the International Conference on Learning Representations (ICLR), 2014.

[24] Sanghee Kang, Younghyun Na, Sung Yup Joung, Sun Il Lee, Sang Cheul Oh, and Byung Wook Min. The significance of microsatellite instability in colorectal cancer after controlling for clinicopathological factors. Medicine, 97(9), 2018.

[25] S Popat, R Hubner, and RS Houlston. Systematic review of microsatellite instability and colorectal cancer prognosis. Journal of clinical oncology, 23(3):609–618, 2005.

[26] Byung-Hoon Min, Jeong Mo Bae, Eui Jin Lee, Hong Suk Yu, Young-Ho Kim, Dong Kyung Chang, Hee Cheol Kim, Cheol Keun Park, Suk-Hee Lee, Kyoung-Mee Kim, et al. The cpg island methylator phenotype may confer a survival benefit in patients with stage ii or iii colorectal carcinomas receiving fluoropyrimidinebased adjuvant chemotherapy. BMC cancer, 11(1):1–10, 2011.

[27] Stacey Shiovitz, Monica M Bertagnolli, Lindsay A Renfro, Eunmi Nam, Nathan R Foster, Slavomir Dzieciatkowski, Yanxin Luo, Victoria Valinluck Lao, Raymond J Monnat Jr, Mary J Emond, et al. Cpg island methylator phenotype is associated with response to adjuvant irinotecan-based therapy for stage iii colon cancer. Gastroenterology, 147(3):637–645, 2014.

[28] Zhenli Diao, Yanxi Han, Yuqing Chen, Rui Zhang, and Jinming Li. The clinical utility of microsatellite instability in colorectal cancer. Critical reviews in oncology/hematology, 157:103171, 2021.

[29] TO Sillo, AD Beggs, DG Morton, and G Middleton. Mechanisms of immunogenicity in colorectal cancer. Journal of British Surgery, 106(10):1283–1297, 2019.

[30] Leland McInnes, John Healy, and James Melville. Umap: Uniform manifold approximation and projection for dimension reduction. arXiv preprint 1802.03426, 2018.

[31] F. Pedregosa, G. Varoquaux, A. Gramfort, V. Michel, B. Thirion, O. Grisel, M. Blondel, P. Prettenhofer, R. Weiss, V. Dubourg, J. Vanderplas, A. Passos, D. Cournapeau, M. Brucher, M. Perrot, and E. Duchesnay. Scikit-learn: Machine learning in Python. Journal of Machine Learning Research, 12:2825–2830, 2011.

[32] Abien Fred Agarap. Deep learning using rectified linear units (relu). arXiv preprint 1803.08375, 2018.

[33] Diederik P Kingma and Jimmy Ba. Adam: A method for stochastic optimization. arXiv preprint 1412.6980, 2014.

